# HSP60 chaperone deficiency disrupts the mitochondrial matrix proteome and dysregulates cholesterol synthesis

**DOI:** 10.1101/2024.01.31.578131

**Authors:** Cagla Cömert, Kasper Kjær-Sørensen, Jakob Hansen, Jasper Carlsen, Jesper Just, Brandon F. Meaney, Elsebet Østergaard, Yonglun Luo, Claus Oxvig, Lisbeth Schmidt-Laursen, Johan Palmfeldt, Paula Fernandez-Guerra, Peter Bross

## Abstract

Mitochondrial proteostasis is critical for cellular function and survival. HSP60 is a molecular chaperone that interacts with more than 260 mitochondrial matrix proteins to assist in their folding and few genetic variants of HSP60 are compatible with life. The few reported human patients with HSP60 variants show phenotypes of neurodevelopmental delay associated with brain hypomyelination. It is currently unknown how deficiency of the HSP60 links to hypomyelination. Here, we studied the onset and progression of HSP60 deficiency in: (1) a HSP60 mutation-inducible cell system, (2) skin fibroblasts from patients with disease-associated HSP60 variants, and (3) zebrafish HSP60 knockout larvae. Collectively, we show how HSP60 deficiency leads to pervasive dysfunctions: (1) downregulated mitochondrial matrix proteome, (2) transcriptional activation of cytosolic stress responses, (3) and lipid accumulation with dysregulated cholesterol biosynthesis. In zebrafish larvae HSP60 deficiency induced early developmental abnormalities. Our comprehensive data identifies HSP60 as a master regulator of mitochondrial proteostasis and suggests a pivotal effect of HSP60 dysfunction on myelination through dysregulation of cholesterol biosynthesis.

**Highlights:**

- HSP60 deficiency disrupts mitochondrial matrix proteins activating stress responses
- Disrupted matrix proteome impairs catabolism causing acetyl-CoA shortage
- HSP60 deficiency dysregulates cytosolic cholesterol synthesis
- Hsp60 deficiency causes developmental abnormalities in zebrafish larvae
- Dysregulated cholesterol synthesis links HSP60 deficiency to hypomyelination

**Graphical abstract:** 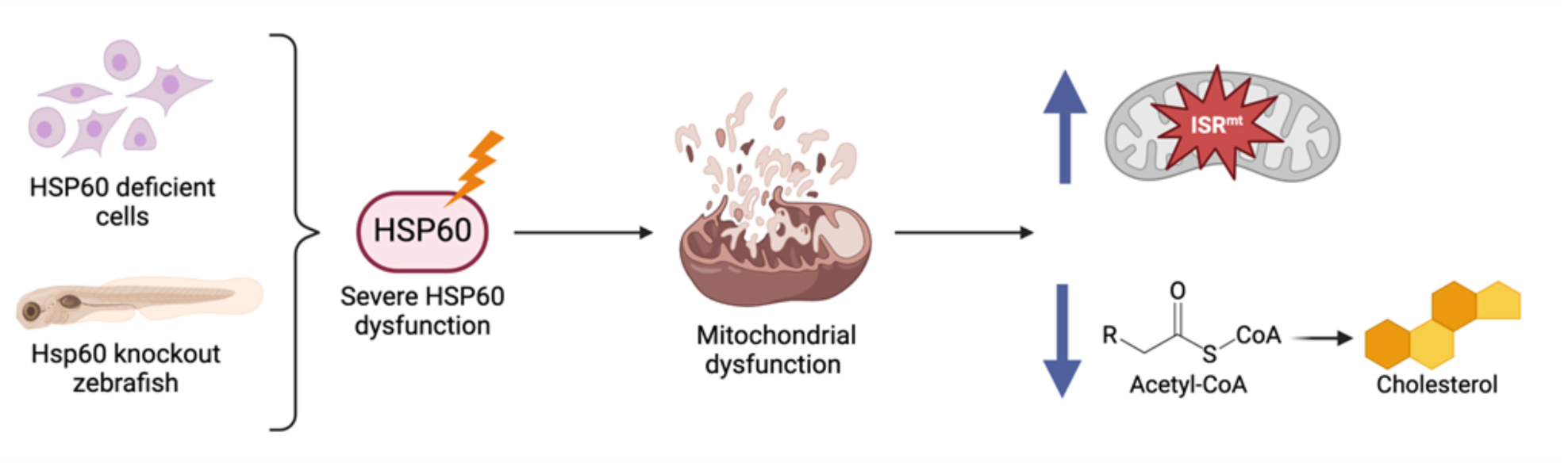

## Introduction

Mitochondria play an indispensable role in the cellular metabolism of eukaryotic cells. In addition to their well-known “powerhouse” function, responsible for the production of the majority of cellular ATP through oxidative phosphorylation, they serve as enzymatic machineries for the production of essential metabolites and possess signaling mechanisms that control the proliferation of eukaryotic cells ^1^. High energy-consuming tissues, such as the central nervous system, which consumes 20% of total body oxygen but accounts for only 2% of total body weight, are highly dependent on mitochondrial functions ^2^. Disturbances of mitochondrial functions are implicated in the pathogenesis of many neurological diseases ^3,4^.

Mitochondria contain approximately 1100 proteins. Most of these proteins are nuclear-encoded, and about half are localized in the mitochondrial matrix space ^5^. Heat Shock Protein 60 (HSP60; gene name *HSPD1*) and Heat Shock Protein 10 (HSP10) are ubiquitously expressed chaperones in the mitochondrial matrix. They assemble into barrel-shaped complexes that encapsulate client proteins and promote correct folding. These chaperones thus form part of the mitochondrial proteostasis system that maintains protein homeostasis ^6,7^. HSP60 interacts with more than 260 mitochondrial proteins ^8^, so its dysfunction potentially affects multiple mitochondrial functions. Because of the broad impact, knocking out HSP60 is lethal in mice and zebrafish ^9,10^, and very few genetic HPS60 defects have been detected in human patients. Patients with disease-associated variations in HSP60 present with neurodevelopmental disorders with impaired myelination as a common feature ^11–15^. Besides *HSPD1* which encodes HSP60, two other genes encoding mitochondrial proteins have been reported to be associated with hypomyelinating leukodystrophies: AIF1, and PYCR2 ^16,17^. Myelination requires local cholesterol synthesis, which depends on metabolites provided by mitochondria ^18,19^. HSP60 deficiency may therefore cause disturbances in mitochondrial metabolisms that feed into the synthesis of cholesterol and other myelin lipids.

Early fetal death of homozygous HSP60 knockout mice ^9^, rapid cell death of knockout tissues in a conditional HSP60 knockout mouse model ^20,21^, and slow onset of phenotypes in heterozygous HSP60 knockout mice ^22^, have hampered studies to monitor pathogenic processes associated with HSP60 deficiency, and in particular, the mechanisms of myelination impairment.

To elucidate the pathogenetic processes, we studied the consequences of various degrees of HSP60 deficiency using three complementary systems: (1) an engineered HEK293 cell line carrying an inducible dominant-negative HSP60 mutant allele (HSP60-D423A) to monitor severe HSP60-deficiency over time; (2) fibroblasts from patients with disease-associated HSP60 variants to study the effects on cells from patients with this ultra-rare disease; and (3) Hsp60 knockout zebrafish larvae to evaluate severe Hsp60-deficiency during early development. We characterized mitochondria-related phenotypes and global transcriptome proteome changes. In this way, we comprehensively mapped the primary effects of HSP60 deficiency in the mitochondrial matrix space, secondary effects on the activation of cellular stress responses, and alterations in cellular metabolism. Our findings extend the understanding of the mechanistic consequences of HSP60 deficiency at the molecular, cellular, and whole organism levels.

## Results

### HSP60 dysfunction results in reduced cell proliferation and progressing perturbations in key mitochondrial functions

To investigate the immediate effect of HSP60 dysfunction on mitochondrial functions, we induced the expression of a dominant-negative ATPase-deficient HSP60 mutant (HSP60-D423A) in HEK293 cells ^23^ using tetracycline **(Figure 1A)**. Incorporation of HSP60-D423A subunits into HSP60 heptamer rings leads to dysfunction of the chaperonin complex. We titrated tetracycline levels to have a biologically relevant cellular model that can be monitored over time. Expression of the HSP60-D432A protein resulted in significantly reduced cell counts at 72 hours of induction compared to uninduced and HSP60-WT co-expressing cells **(Figure 1B).** Cell viability was not significantly affected **(Figure S1A)**. There was no growth difference between cells induced for the WT transgene and uninduced cells. We subsequently studied the consequences of HSP60-D423A expression on mitochondrial functions within 72 hours of induction. Very low levels of leaky expression of the mutant HSP60 transcript were detected by RNA sequencing in uninduced cells **(Figure 1C)**. The fraction of HSP60-D432A transcript and protein levels increased strongly upon tetracycline induction accounting for approximately 80% of total HSP60 transcript and approximately 50% of total HSP60 protein **(Figures 1C and 1D)**. Total HSP60 transcript levels increased more than 2-fold, but total HSP60 protein levels increased by only about 40% after 48 hours of induction and were maintained for 72 hours.

**Figure 1:**
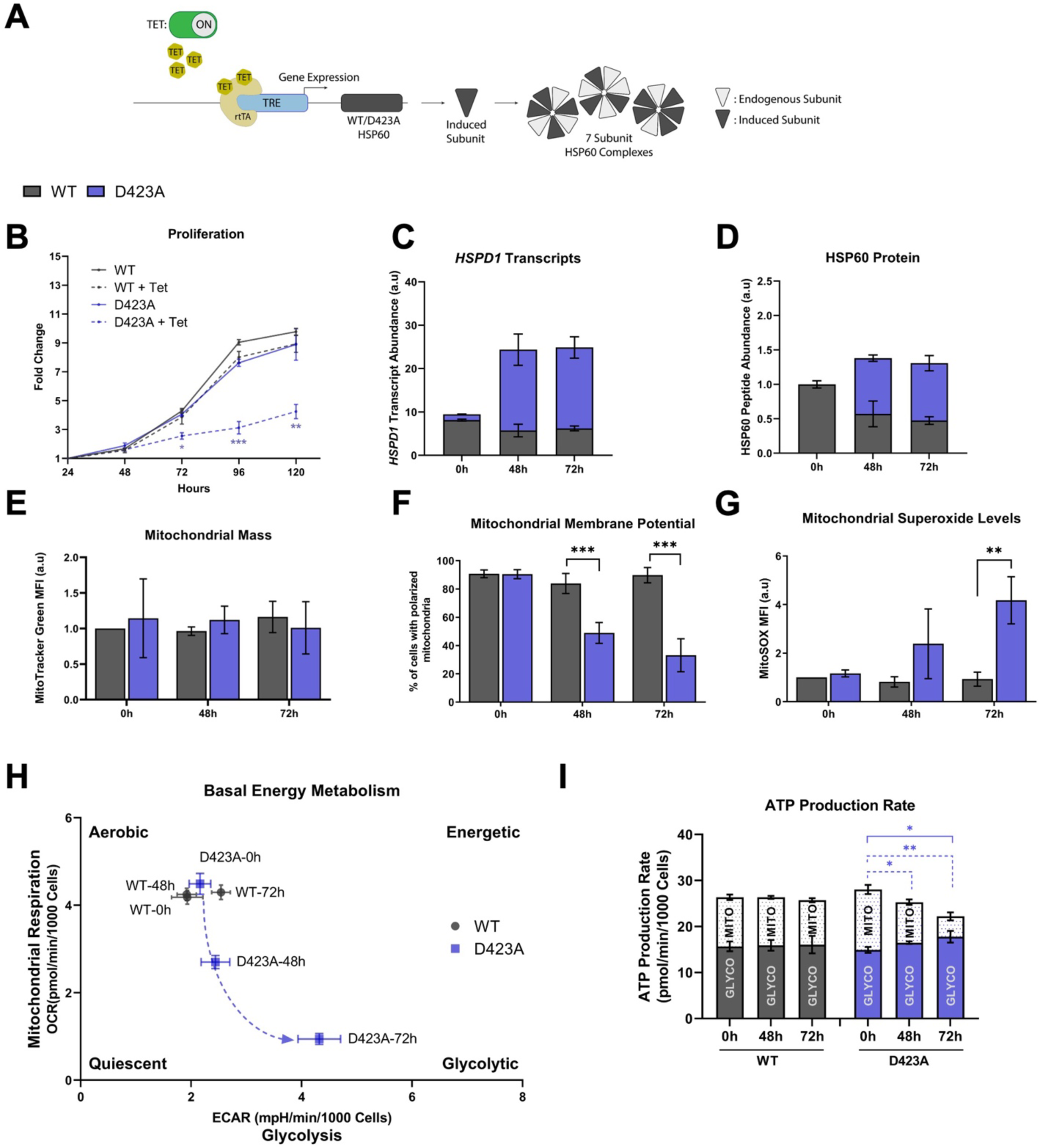
HSP60 dysfunction results in cell proliferation arrest and progressing perturbations in key mitochondrial functions. **(A)** Schematic representation depicting the mechanism in tetracycline-inducible HEK293 cell lines expressing wild-type HSP60 (WT) or the HSP60 dominant-negative ATPase-deficient mutant HSP60 (D423A). Inducible expressed subunits integrate with endogenously expressed subunits to form heteromeric seven-subunit HSP60 complexes. **(B)** Cell proliferation of WT and D423A cell lines, uninduced and induced with 50 ng/mL tetracycline for different time intervals. Represented in fold change compared to initial cell counts (statistical comparison between WT and D423A induced cell lines). **(C-D)** The abundance of D423A and endogenous WT *HSPD1* transcript **(C)** and HSP60 protein **(D)** levels, quantified by RNA sequencing and mass spectrometry, respectively, at 0h (uninduced), 48h, and 72h induced D423A cells. **(E)** Quantification of mitochondrial mass by MitoTracker green fluorescence in induced WT and D423A cell lines (n≥3) at 0h (uninduced), 48h, and 72h induced D423A cells. **(F)** Percentage of cells with intact mitochondrial membrane potential in induced WT and D423A cells by JC-1 fluorescence (n≥3) at 0h (uninduced), 48h, and 72h induced D423A cells. **(G)** Mitochondrial superoxide levels in induced WT and D423A cells, based on MitoSOX fluorescence (n≥3) at 0h (uninduced), 48h, and 72h induced D423A cells. **(H)** Bioenergetic profiles of induced WT and D423A cells in basal conditions, analyzed with Seahorse XFe96 analyzer at 0h (uninduced), 48h, and 72h induced D423A cells. **(I)** Total ATP production distinguishing the proportion of mitochondrial and glycolytic ATP production rates, analyzed using the Seahorse XFe96 analyzer at 0h (uninduced), 48h, and 72h induced D423A cells. Blue continuous line compares the total ATP production rate, and blue dashed lines compare the mitochondrial ATP production rate. Graphs represent mean ±SD (n=3); statistical significance was determined with multiple unpaired t-tests with Holm-Sidak correction between WT and D423A induced cell lines unless otherwise stated (* p < 0.05, ** p < 0.01, *** p < 0.001). Abbreviations: ECAR, extracellular acidification rate (represents glycolysis); FCCP, Carbonyl cyanide 4-(trifluoromethoxy) phenylhydrazone; h, hours; HEK, human embryonic kidney; HSP, heat-shock protein; OCR, oxygen consumption rate (represents oxidative phosphorylation); SD, standard deviation; TET, tetracycline; tTA, reverse tetracycline transactivator; TRE, tetracycline-responsive promoter element; WT, wild-type.

The induction of HSP60-D423A did not lead to significant changes in mitochondrial mass compared to the induction of HSP60-WT **(Figure 1E)**. However, key mitochondrial functions were affected in a time-dependent manner, including decreased mitochondrial membrane potential and increased mitochondrial superoxide levels **(Figures 1F and 1G)**. Moreover, mitochondrial respiration and mitochondrial ATP production rates were decreased in the HSP60-D432A induced cells without changing the cellular ADP/ATP ratio. **(Figures 1H and 1I; Figures S1B-S1D)**. The decrease in mitochondrial ATP production was partially compensated by an increase in glycolytic ATP production at 48 hours **(Figures 1I; Figure S1E)**. However, the respiration defect in the HSP60-D432A induced cells was too severe to be compensated at 72 hours, where the total ATP production rate was significantly decreased **(Figure 1I; Figure S1F)**.

### HSP60 dysfunction severely affects the mitochondrial proteome

Our previous studies using cellular knock-down and heterozygous knock-out mouse models ^24,25^ had shown that HSP60 deficiency leads to impaired folding of certain mitochondrial matrix proteins and hence their increased degradation. Based on this, we hypothesized that HSP60 dysfunction would lead to decreased levels of those mitochondrial proteins that depend on HSP60 for proper folding. In parallel, the levels of the encoding transcripts would remain unchanged or even increased due to compensatory mechanisms. In order to test this hypothesis, we quantified the proteomes and transcriptomes of uninduced, 48h, and 72h induced D423A cells **(Figure S2A)**. Unsupervised principal component analysis of proteomics and transcriptomics results showed that uninduced, 48h, and 72h samples clustered separately **(Figures S2B and S2C)**. We quantified a total of 15,989 genes at RNA level and 6,359 proteins. Compared to uninduced D432A cells 112 (48h) and 566 (72h) genes (FDR p<0.05 and |FC|>2) were differentially expressed at the RNA level, and 258 (48h) and 485 (72h) proteins at the protein level (p<0.05 and |FC|>1.2) **(Figure 2A and 2B)**. At the RNA level, differentially expressed genes were mainly upregulated at both 48h (63 genes, 82%) and 72h (435 genes, 70%), and only a few of the upregulated genes encoded mitochondrial proteins **(Figures 2C and 2E)**. The increased number of upregulated genes at 72h compared to 48h suggests the activation of compensatory mechanisms (see enrichment analysis section). In contrast, the majority of differentially expressed proteins were downregulated at both time points, 48h (215 proteins, 92%) and 72h, (385 proteins, 86%). In line with our hypothesis, most of the downregulated proteins represented mitochondrial proteins (distinguished according to MitoCarta 3.0 ^5^) **(Figures 2D and 2F; Figure S2D).**

**Figure 2:**
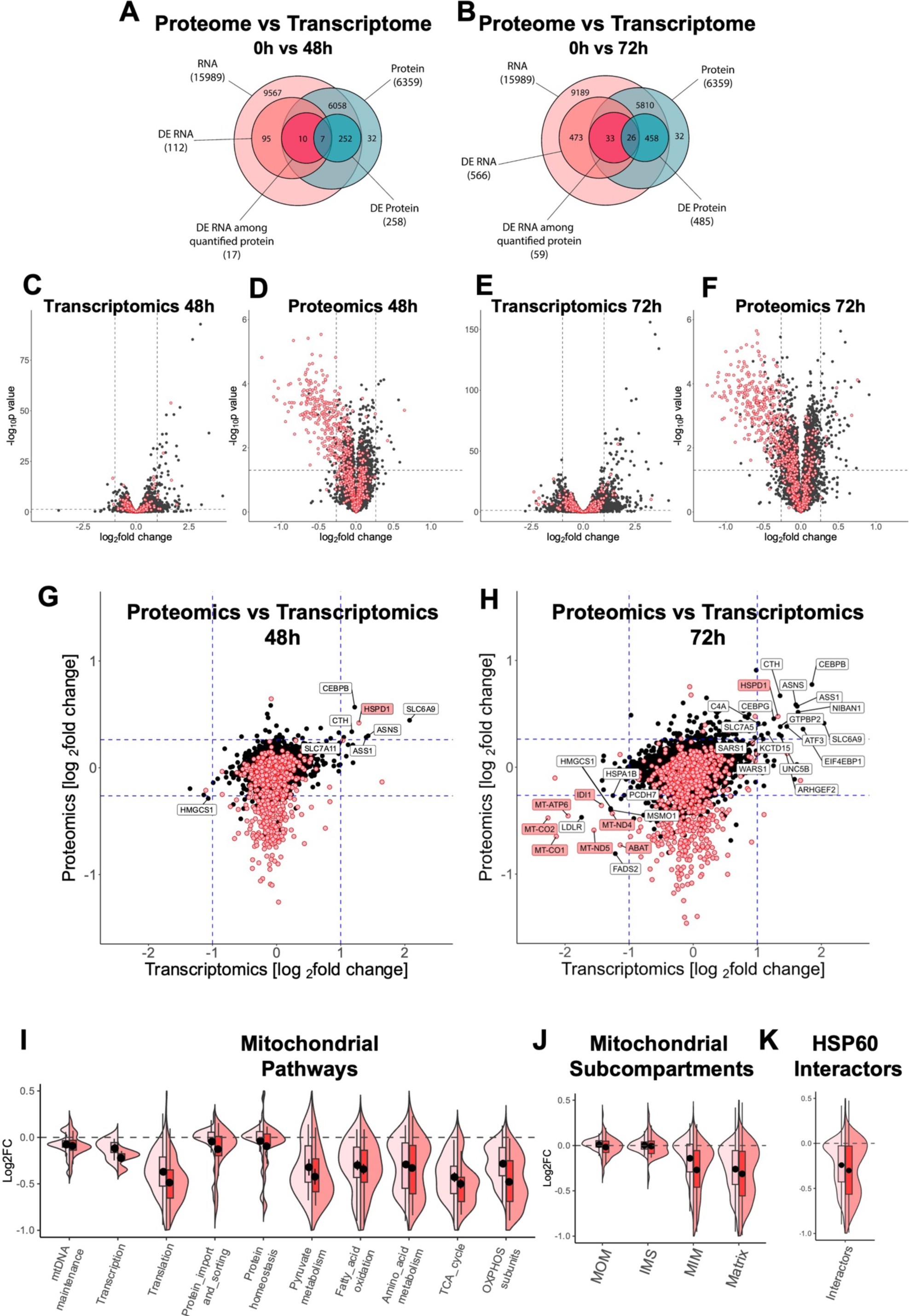
HSP60 dysfunction severely regulates the mitochondrial proteome, but not the transcriptome. **(A-D)** Volcano plots showing the comparison of induced D423A cells at 48h and 72h with 0h for quantified transcriptome **(A, C)**, and proteome **(B, D)**. Cut-off values are indicated with dashed lines. Mitochondrial proteins are indicated in red. **(E-F)** Venn diagrams representing the number of quantified and differentially expressed genes in transcriptomics and proteomics. D423A cells at 0h were compared to 48h **(E)** or 72h induced D423A cells **(F)**. The pairing of quantified genes of transcriptomics and proteomics was performed through Ensembl gene IDs. **(G, H)** Comparison plot of log2fold changes of Proteomics versus Transcriptomics at 48h **(G)** and 72h **(H)**. Cut-off values are indicated with dashed lines. Mitochondrial proteins are highlighted in red. Genes significantly up- or down-regulated at both transcript and protein level are labeled. **(I-K)** Violin plots distinguishing proteins from mitochondrial pathways **(I)**, mitochondrial sub-compartments **(J)**, and HSP60 interactors **(K),** and of induced D423A cells at 0h in comparison to 48h (light red) or 72h induced D423A cells (dark red). Cut off values for transcriptomics: Benjamini-Hochberg adjusted *p*-value <0.05 and |log2FC|>1. Cut off values for proteomics: *p-*value <0.05 and |log2FC|>0.26 Abbreviations: DE, differentially expressed; FC, fold change; IMS, intermembrane space; MIM, mitochondrial inner membrane; MOM, mitochondrial outer membrane (See also the legend of Figure 1).

Plots depicting fold changes of proteomics versus transcriptomics clearly illustrate that for the majority of mitochondrial proteins, downregulation was not due to transcriptional downregulation **(Figures 2G and 2H)**. Seven of the 12 genes significantly downregulated at both RNA and protein levels at 72h induction encode mitochondrial proteins (labels highlighted in red). Five of them are encoded by the mitochondrial DNA (mtDNA) and they are likely decreased because proteins involved in mtDNA transcription and translation are compromised. Of note, we observed significant upregulation of genes involved in the mitochondrial integrated stress response (ISR^mt^) (ATF3, CTH, ASNS) and downregulation of genes concerned with cholesterol synthesis (HMGCS1, MSMO1) and lipid transport (LDLR) **(Figure 2H)**. A more detailed discussion is provided in the enrichment analysis section.

The fold change distribution of mitochondrial proteins revealed that many of the downregulated proteins are part of mitochondrial pyruvate metabolism, fatty acid oxidation, amino acid degradation, TCA cycle, OXPHOS, and mtDNA translation **(Figure 2I)**. Proteins involved in the maintenance and transcription of mtDNA, representatives of the mitochondrial protein import and sorting machinery, and components of the mitochondrial protein homeostasis system were only slightly affected, even at 72h. Violin plots of an extended number of mitochondrial pathway groups further support the notion that especially functions in the matrix and inner membrane were extensively affected by HSP60 deficiency whereas proteins in the other mitochondrial compartments were not affected **(Figure S2E)**. In larger complexes, such as respiratory chain complex I or mitochondrial ribosomes, the effect of compromising several subunits appears to destabilise the whole complex. When the fold change distributions of the proteins were differentiated according to the mitochondrial subcompartments in which they are localized, mainly matrix and inner membrane proteins were downregulated **(Figure 2J)**.

We also comparatively analyzed the fold change distributions of the proteins interacting with HSP60 ^8^. Mitochondrial interactors showed a bimodal distribution that was most evident at 72 hours of induction, where two separate peaks are clearly distinguished **(Figure 2K)**. This suggests that for a fraction, but not all the interactors, interaction with HSP60 is essential for folding. Similar bimodal distributions with two distinct peaks were also seen in the violin plots for matrix proteins and metabolism groups **(Figures 2I and 2J)**. The observation that a considerable fraction of matrix proteins and interactors were almost unchanged in amounts suggests that import into the matrix was not significantly impaired. Taken together, our results suggest that HSP60 dysfunction leads to severe disruption of the mitochondrial matrix proteome and associated functions.

### Patient fibroblasts carrying HSP60 disease-associated variants present mild mitochondrial dysfunction with cytosolic compensatory responses

The D423A-HSP60 cell system represents the effects of a very severe ATPase domain mutation that is not reported in patients with disease-associated variants and is also absent in the Genome Aggregation Database (gnomAD ^26^). To evaluate whether disease-associated variations found in patients trigger similar changes in the mitochondrial proteome, we performed proteomics analysis of dermal fibroblasts from two patients carrying disease-associated *HSPD1* variants and presenting with hypomyelination phenotype. Patient 1 (P1) is heterozygous for the *de novo* variation p.Leu47Val with the milder disease phenotype ^14^ and Patient 2 (P2) is homozygous for the p.Asp29Gly variant causing the severe lethal hypomyelinating leukodystrophy MitCHAP60 disease ^12^. Functional assays have shown that both HSP60 variants possess residual chaperone activity ^14,15^. In contrast to our D423A-HSP60 cell system, the proteomes of both patient cell lines, each compared to three healthy control individuals, showed mainly upregulation of differentially expressed proteins **(Figures 3A and 3B)**. Only a few mitochondrial proteins were significantly regulated in the two patients; none was downregulated in P2 and only two in P1: MGST1, a glutathione-S transferase, and IARS2, the mitochondrial t-RNA ligase for isoleucine. The up-regulated mitochondrial proteins comprised peroxidases (PRX4 and PRDX5), outer membrane proteins (MITCH1 and NIPSNAP2), and fatty acid oxidation enzymes (ECH1, ACAA2). Consequently, fold change distribution analysis, distinguishing according to mitochondrial subcompartments, showed a tendency to decreased levels of matrix proteins only in P1 **(Figure 3C)**.

**Figure 3:**
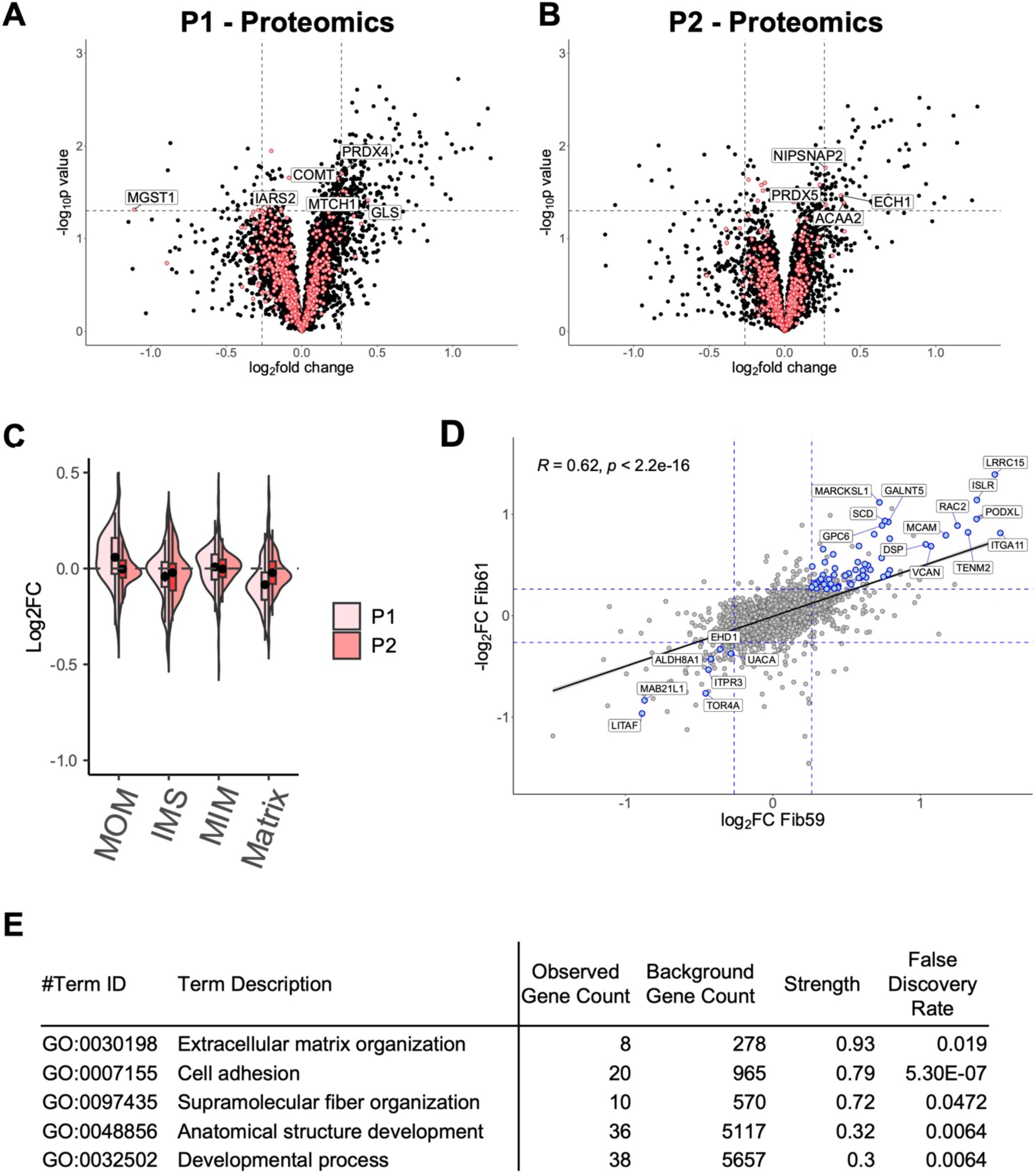
Proteomics analysis of patient fibroblasts carrying HSP60 disease-associated variants. **(A-B)** Volcano plots showing the quantified proteome of human dermal fibroblasts of the Patient 1 (heterozygous HSP60-p.Leu47Val) **(A)** and of human dermal fibroblasts of the Patient 2 (homozygous HSP60-p.Asp29Gly) **(B)** compared to their respective human dermal fibroblast control cells (n = 3). Cut-off values are indicated with dashed lines (*p-* value <0.05 and |log2FC|>0.26). Mitochondrial proteins are indicated in red. **(C)** Violin plot distinguishing the protein Log2fold changes from dermal fibroblasts from Patient 1 (light red) and Patient 2 (dark red) in comparison to their respective controls. **(D)** Correlation of the protein Log2fold changes of Patient 1 (horizontal axis) with Patient 2 (vertical axis) in comparison to their respective controls. The correlation line is based on a linear mode; Pearson R and *p* value are given. **(E)** Enriched GO terms for significantly up-regulated fibroblast proteins in patient 1 and patient 2 as analyzed using the STRING database (https://string-db.org).

The overall fold changes of the protein levels of the two patients were correlated (R = 0.62) (**Figure 3D**), suggesting that similar responses were activated in cells from both patients. Considering only the fold changes of proteins with a *p*-value <0.05 in the triplicate analyses for both patients, a significant upregulation of a larger group of non-mitochondrial proteins in both patient samples was observed, while only seven proteins were significantly downregulated. Interaction network analysis of the significantly upregulated proteins revealed clusters of proteins associated with extracellular functions and fiber organization **(Figure 3E)**.

### Hsp60 knockout triggers phenotypes indicating developmental abnormalities in zebrafish larvae

To understand the effects of Hsp60 deficiency in developing vertebrates, we generated CRISPR/Cas9-mediated *hspd1* knockout zebrafish lines by targeting exon 2 to induce a frameshift mutation. We selected an allele with a 56 base pair deletion inducing a frameshift mutation leading to loss of protein functions **(Figures S3A and S3B)**. The lack of the Hsp60 protein in the *hspd1* knockout *(hspd1^-/-^)* embryos at day 3 post fertilization (3 dpf) and at early larval stages (5 and 7 dpf) confirmed the efficiency of the Hsp60 protein knockout **(Figure 4A)**. The truncated *hspd1* knockout transcript was present in *hspd1*^-/-^ larvae, but in lower amounts than the full-length wild-type transcript in wild-type *(hspd1^+/+^)* larvae **(Figure 4C)**. Morphologic inspection indicated that *hspd1^-/-^* larvae displayed a mildly dysmorphic phenotype compared to the *hspd1^+/+^,* while heterozygous embryos *(hspd1*^+/-^) did not **(Figure 4B)**. Specifically, *hspd1*^-/-^ larvae had a shorter body length at 5 dpf, and a smaller eye area at 5 and 7 dpf (**Figures 4D and 4E; Figure S3C).** The majority of knockout larvae failed to inflate the swim bladder at 5 and 7 dpf **(Figure 4F)**. No significant differences in these parameters were observed at 3 dpf. Overall, this indicated that Hsp60 deficiency causes early larval developmental abnormalities.

**Figure 4:**
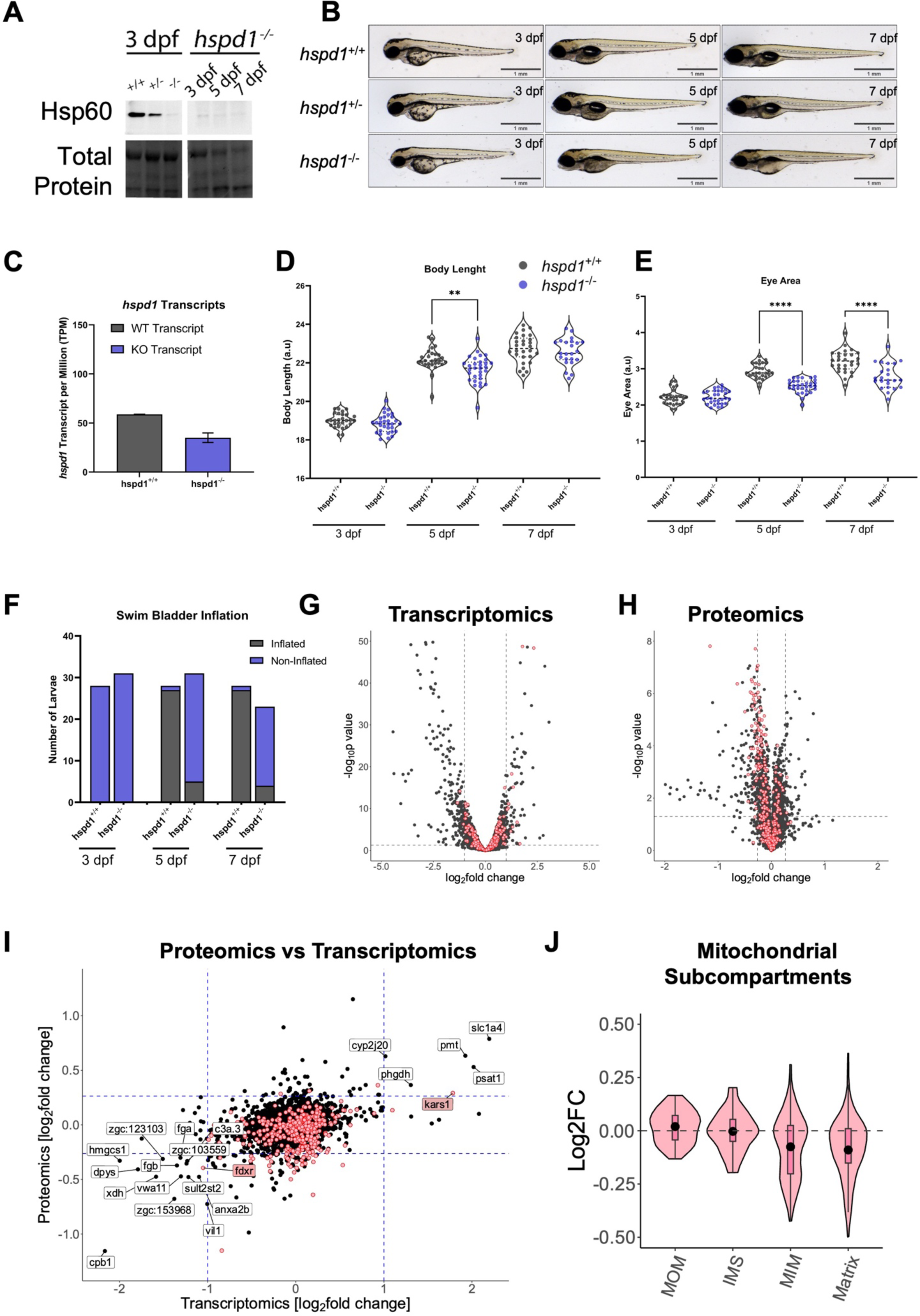
Zebrafish Hsp60 knockout results in early larval developmental abnormalities. **(A)** Hsp60 protein levels of *hspd1^+/+^*, *hspd1^+/-^*, and *hspd1^-/-^* larvae at 3 dpf, and *hspd1^-/-^* larvae at 3, 5, and 7 dpf. Assessed by western blot (n=6). **(B)** Lateral imaging of *hspd1^-/-^* larvae at 3, 5, and 7 dpf indicated developmental delay at 5 and 7 dpf, but *hspd1+/-* larvae showed no significant phenotypic differences compared to *hspd1^+/+^* larvae. **(C)** Quantification of WT and KO *hspd1* transcript levels in *hspd1^+/+^* and *hspd1^-/-^* larvae at 5 dpf, quantified by RNA sequencing. **(D-F)** Phenotypic quantification of body length **(D)**, eye size **(E)**, and swim bladder inflation **(F)** in *hspd1^+/+^* (n = 28), *hspd1^-/-^* (n = 31) larvae at 3, 5, and 7 dpf. **(G-H)** Volcano plots showing analysis of transcriptome **(G)**, and proteome **(H)** changes between *hspd1^+/+^* and *hspd1^-/-^* larvae. Cut-off values are indicated with dashed lines. Mitochondrial proteins were indicated in red. **(I)** Comparison plot of log2fold changes of Proteomics versus Transcriptomics in *hspd1^-/-^* larvae. Cut-off values are indicated with dashed lines. Mitochondrial proteins are highlighted in red. Genes significantly up- or down-regulated at both transcript and protein level are labeled. **(J)** Violin plots distinguishing proteins from mitochondrial sub-compartments of *hspd1^-/-^* larvae in comparison to *hspd1^+/+^* larvae. Statistical significance was determined with one-way ANOVA: * p < 0.05, **** p < 0.0001. Abbreviations: dpf, days post-fertilization; KO, knockout; sgRNA, single guide RNAs; WT, wild type.

### Hsp60 deficiency results in mitochondrial proteome changes in zebrafish larvae

To further understand the effects of HSP60 deficiency on early larval development, we examined the proteome and transcriptome changes in zebrafish larvae at 5 dpf (**Figure S4A)**. The *hspd1^+/+^* and *hspd1^-/-^* samples clustered separately in the unsupervised principal component analysis of both the proteome and transcriptome data **(Figures S4B and S4C).** A major part of differentially regulated genes (208 genes, 71%) and proteins (179 proteins, 76%) were downregulated. Similar to the D423A-HSP60 cell system, many genes encoding mitochondrial proteins were downregulated at protein level in *hspd1^-/-^* larvae, whereas they were unchanged at RNA level **(Figures 4G and 4H)**. Genes that were significantly up- or downregulated at both the transcript and protein level encoded mainly non-mitochondrial proteins, with only two exceptions (*kars1* and *fdxr*) **(Figure 4I)**. Strikingly, we found that, as in the D432A-HSP60 cell system, ISR^mt^ genes were among the consistently upregulated genes (*psat1*, *phgdh*) and the gene that catalyzes the first step in the mevalonate/cholesterol synthesis pathway (*hmgcs1*) was among the consistently downregulated genes **(Figure 4I)**.

In accordance with the findings in the D423A-HSP60 cell system, analysis of submitochondrial localization revealed that fractions of matrix and inner membrane proteins were downregulated in *hspd1^-/-^* larvae, whereas outer membrane and intermembrane space proteins were unaffected **(Figure 4J)**. Comparison of the distributions of fold changes of mitochondrial proteins in *hspd1^-/-^* larvae, compared to *hspd1^+/+^* larvae and the D423A-HSP60 cells at 72h induction, compared to uninduced cells, revealed significant correlations for matrix and inner membrane proteins **(Figure S4D)**. This suggests that a fraction of mitochondrial protein orthologs are sensitive to HSP60 deficiency in both species while others are not. In conclusion, as in the HEK D423A-HSP60 cell system, there was a clear reduction in amounts of many mitochondrial matrix and inner membrane proteins in the zebrafish HSP60 knockout larvae.

### Enrichment analysis reveals affected fatty acid and cholesterol metabolism

To elucidate adaptive and compensatory mechanisms caused by HSP60 deficiency, we performed enrichment analysis for the differentially expressed proteins and genes in the D423A-HSP60 cell system and zebrafish HSP60 knockout studied, using Enrichr and FishEnrichr ^27,28^. Not surprisingly, the enriched clusters from proteomics in both systems were overwhelmingly related to mitochondrial functions and pathways, such as mtDNA translation, oxidative phosphorylation (OXPHOS), fatty acid oxidation (FAO), and TCA cycle **(Figure 5A and 5B)**. Contrarily, transcriptomics-enriched clusters showed few mitochondria-related terms. One of the common cellular responses to mitochondrial dysfunction is the activation of the ISR^mt^, allowing cells to adapt to unfavorable conditions. Both the induced HEK D423-HSP60 cells and the zebrafish HSP60 knockout larvae presented significant upregulation of many genes of the ISR^mt^ at both RNA (**Figures 5C and 5D)** and protein levels **(Figures S5A and S5B)**. However, the mitochondrial ISR^mt^ proteins SHMT2, CLPP, and MTHFDH2 were decreased in the HEK D423-HSP60 cell system, whereas the encoding transcripts for MTHFD2 and SHMT2 were upregulated **(Figures 5C and S5A).** This suggests that the mitochondrial branch of the ISR^mt^ response could not be executed at the protein level. Respiratory chain deficiency leads to an upregulation of metabolites from the one-carbon cycle branch of the ISR^mt^, e.g. glycine, serine, and threonine ^29^. Consistent with this, targeted metabolite analysis showed increased levels of the amino acids glycine, serine, and threonine in the D423A-HSP60 cell system and serine and glycine in the *hspd1^-/-^* larvae **(Figures 5E and 5F)**.

**Figure 5:**
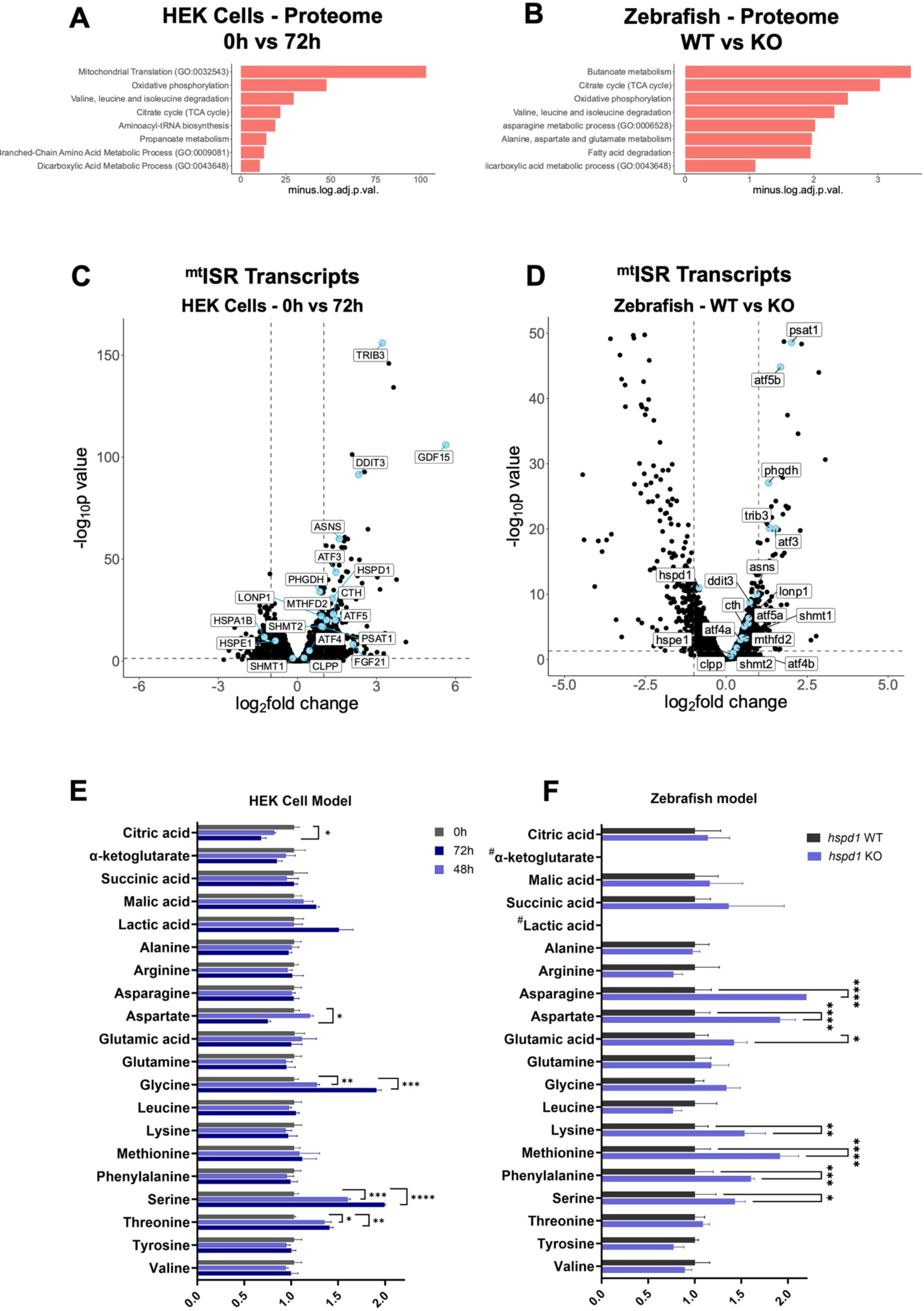
ISR^mt^ is upregulated in D423A cells and *hspd^-/-^* larvae. **(A-B)** Enrichment analysis of differentially expressed proteins of D423A cells at 0h in comparison to 72h induced D423A cells **(A)** and in 5 dpf *hspd1^-/-^* larvae in comparison to *hspd1^+/+^* larvae **(B)**. Performed on EnrichR at gene name level for proteomics GO biological process and KEGG terms are included. The enriched terms are shown as a negative log10 of adjusted *p*-values. Adjusted *p*-values <0.05 were considered significant. **(C-D)** Volcano plots showing the transcriptomic regulation of mitochondrial integrated stress response (ISR^mt^) transcripts in D423A cells at 0h in comparison to 72h induction **(C)** and in 5 dpf *hspd1^-/-^* larvae in comparison to *hspd1^+/+^* larvae **(D)**. **(E-F)** Amino acids and TCA cycle intermediate metabolites were analyzed by targeted metabolomics, and mean-centered abundance values were further normalized to total ion intensity. **(E)** In the D423A cells, significance was calculated against uninduced cells (mean ±SD, n=3, multiple unpaired t-tests with Holm-Sidak correction). **(F)** 5 dpf *hspd1^-/-^* larvae in comparison to *hspd1^+/+^* larvae (mean ±SD, n=3, two-way ANOVA with Sidak’s multiple comparisons correction). Abbreviations: GO, gene ontology; KEGG, Kyoto Encyclopedia of Genes and Genomes; See also Figures 1 and 4.

Analysis of the transcriptomic changes in both the D423-HSP60 cell system and the zebrafish Hsp60 knockout larvae revealed enrichment of GO and KEGG terms centered on sterol/cholesterol/steroid biosynthesis, immune response pathways, and stress responses **(Figures 6A and 6B)**. All these processes mainly occur outside the mitochondria. In both the D423-HSP60 cell system and the zebrafish Hsp60 knockout larvae, many genes encoding enzymes of the mevalonate and steroid synthesis pathways were downregulated both at RNA **(Figures 6C and 6D)** and protein levels **(Figures S5E and S5F**). In addition, targeted metabolite analysis showed reduced citrate levels in the D423-HSP60 cell system **(Figure 5E)**. Citrate is generated from acetyl-CoA supplied from fatty acid oxidation, pyruvate metabolism, and amino acid degradation in the mitochondria and feeds lipid synthesis outside mitochondria ^30^. All these pathways are impaired in our models of HSP60 deficiency. However, no significant changes in citrate levels were seen in the zebrafish Hsp60 knockout larvae **(Figure 5F)**.

**Figure 6:**
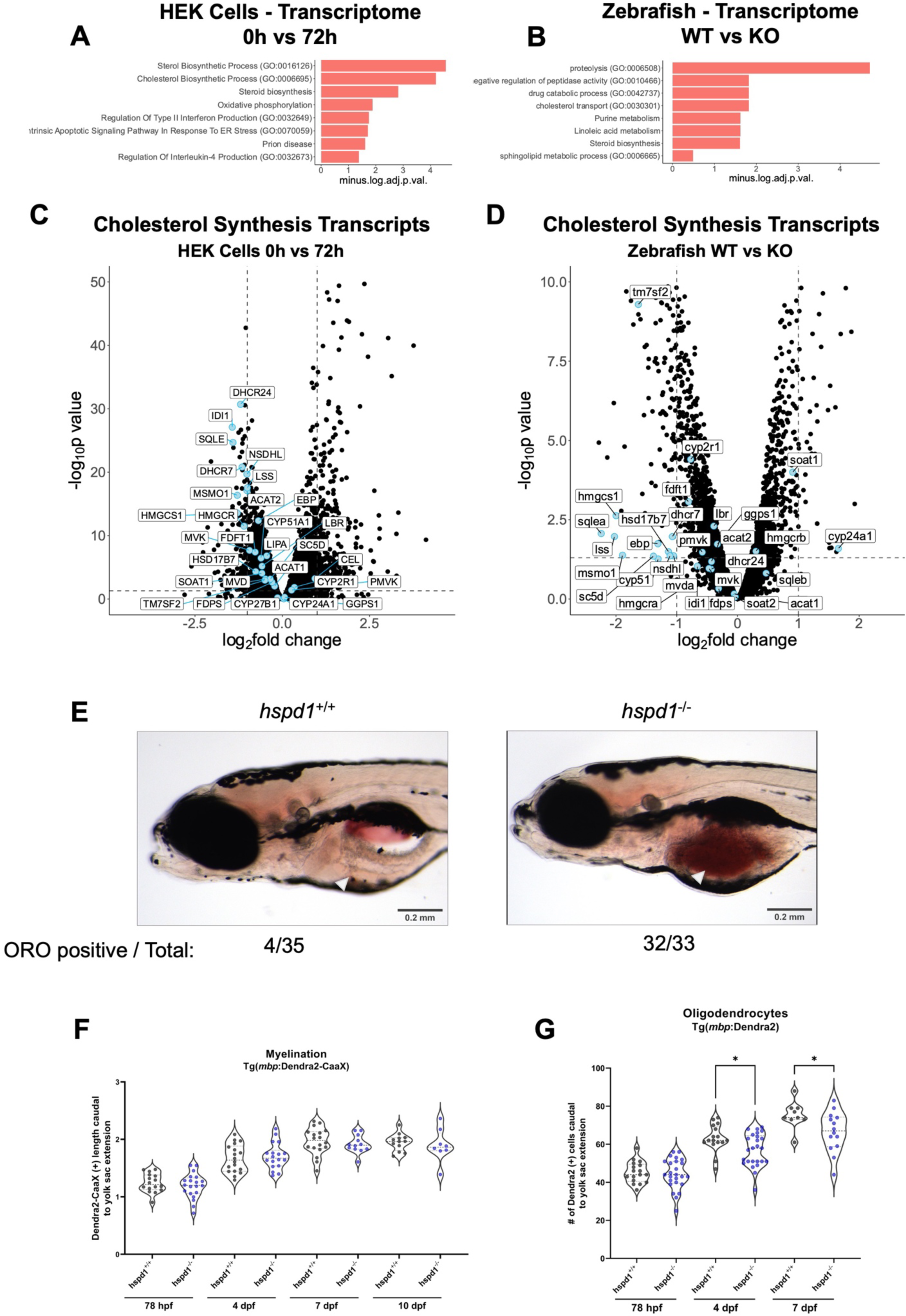
Zebrafish Hsp60 knockout larvae have altered lipid metabolism and oligodendrocyte counts. **(A-B)** Enrichment analysis of differentially expressed genes of D423A cells at 0h in comparison to 72h **(A)** and in 5 dpf *hspd1^-/-^* larvae in comparison to *hspd1^+/+^* larvae **(B)**. Performed on EnrichR at gene name level for proteomics and transcriptomics, GO biological process and KEGG terms are included. The enriched terms are shown as a negative log10 of adjusted *p*-values. Adjusted *p*-values <0.05 were considered significant. **(C-D)** Volcano plots showing the transcriptomic regulation of transcripts involved in cholesterol synthesis pathways in D423A cells induced for 72h **(C)** and in 5 dpf *hspd1^-/-^* larvae **(D)**. **(E)** Whole-mount Oil Red O staining of 8 dpf zebrafish larvae shows lipid accumulation and altered lipid composition in the yolk of *hspd1*^-/-^ larvae compared to *hspd1^+/+^* larvae. **(F)** The length of Dendra2-CaaX(+) signal was measured caudal to yolk sac extension at 78 hpf, 4 dpf, 7 dpf, and 10 dpf. Larvae were generated from *hspd1^+/-^ to* Tg(*mbp*:Dendra2-CaaX) mating. **(G)** The number of mature oligodendrocytes was quantified as caudal to yolk sac extension Dendra2-positive cells at 78 hpf, 4 dpf, and 7 dpf. Larvae were generated from *hspd1^+/-^ to* Tg(*mbp*:Dendra2) mating. Abbreviations: TCA, tricarboxylic acid; GO, gene ontology; KEGG, Kyoto Encyclopedia of Genes and GenomesSee also Figure 1, 2 and 4.

These results suggest that severe HSP60 deficiency initially causes dysfunction in many mitochondrial metabolic and energy-producing pathways triggering transcriptional regulations of extramitochondrial pathways. Notably, in the fibroblasts from P1, we saw a significant downregulation of ISR^mt^ proteins and an upregulation of cholesterol synthesis enzymes; both changes were in the opposite direction compared to the HEK and zebrafish models **(Figures S5C, S5D, S5G and S5H)**.

### Hsp60 knockout zebrafish larvae display altered lipid metabolism and a reduced number of oligodendrocytes

As cholesterol is a highly abundant component of myelin, the combination of a shortage of the building block acetyl-CoA and downregulation of the cholesterol synthesis pathway may be the underlying cause of the hypomyelination phenotype observed in patients with HSP60 deficiency. To address this, we assessed the lipid content of whole-mount zebrafish larvae at 8 dpf using Oil Red O staining. In line with a decreased capacity of the fatty acid degradation machinery, the staining showed retention of neutral lipids in the yolk/liver region of *hspd1*^-/-^ larvae at 8dpf **(Figure 6E).** We then investigated myelination in the *hspd1^-/-^* larvae by monitoring myelin sheaths using the transgenic line: Tg(*mbpa*:Dendra2-CaaX). No significant difference in the progression of myelination between knockout and wildtype larvae at 4, 7, and 10 dpf was observed **(Figure 6F; Figure S6A).** Consistent with unaltered myelination, myelin proteolipid protein (Plp) levels were similar in *hspd1*^-/-^, *hspd1*^+/-^, and *hspd1*^+/+^ larvae at 3 dpf, and Plp protein levels in *hspd1*^-/-^ larvae were increased at 5 and 7 dpf compared to 3 dpf **(Figure S6B)**. When using the transgenic line Tg(*mbpa*:Dendra2) with fluorescent labeling of oligodendrocytes, we observed a significantly decreased number of mature oligodendrocytes in the *hspd1*^-/-^ larvae compared to *hspd1*^+/+^ at 4 and 7 dpf **(Figure 6G; Figure S6C)**.

In conclusion, Hsp60 deficiency led to a broad mitochondrial dysfunction and leading to an activation of the ISR^mt^, and a dysfunctional lipid metabolism including dysregulated cholesterol biosynthesis ending in developmental abnormalities during early zebrafish larval stages (**Figure 7)**.

**Figure 7:**
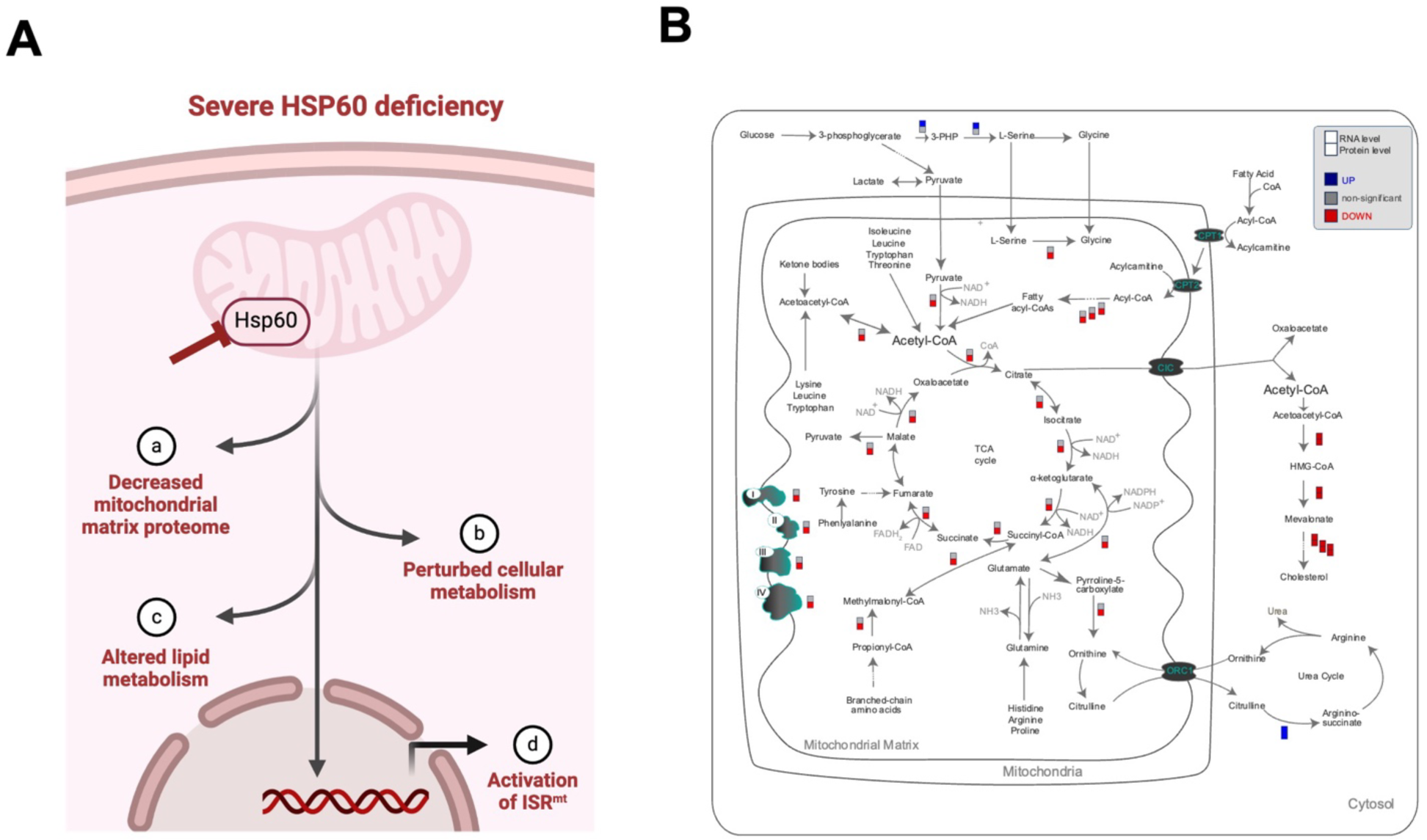
Overview of effects of HSP60 dysfunction. **(A)** Schematic summary of consequences of HSP60 deficiency: (a) mitochondrial matrix protein levels are decreased, (b) leading to changes in cellular metabolism, (c) altering the pool of acetyl-CoA that alters the lipid metabolism (d), and finally activating ISR^mt^. **(B)** Simplified overview summarizing the changes in gene expression of metabolic enzymes and respiratory chain complexes at RNA and protein level affecting metabolic pathways and the respiratory chain as observed in HEK D423A cells induced for 72 h compared to uninduced cells.

## Discussion

In the present study, we performed a comprehensive analysis of cellular and zebrafish models to elucidate the mechanisms and consequences of HSP60 deficiency. **Figure 7A** shows a mechanistic model for the responses on different levels of HSP60 deficiency and **Figure 7B** depicts the major gene expression changes in metabolic pathways observed in the HEK cells expressing the HSP60-D423A variant.

Using a cellular model in which the chaperone function of HSP60 is acutely blocked by the expression of a dominant-negative HSP60 mutant (D423A), we show that a large fraction of the mitochondrial matrix and inner membrane proteins were decreased in abundance. Proteins of all major pathways and functions in the mitochondrial matrix and inner membrane were affected, including the OXPHOS system, amino acid and fatty acid catabolism, and mitochondrial ribosomes **(Figure 2J; Figure S2E)**. Notably, a fraction of mitochondrial HSP60 interactors ^8^ were not or only slightly affected, suggesting that some interacting proteins do not require the help of HSP60 for folding. Levels of mitochondrial proteins that use the TIM22 system to enter the inner membrane, e.g. SLC25-family metabolite carrier proteins ^31^, were stable, suggesting that HSP60 deficiency does not cause a general decrease in the import of mitochondrial proteins. In addition, a considerable fraction of matrix proteins and interactors are not significantly changed in levels suggesting that the decrease of matrix and inner membrane proteins was not due to impaired import.

Similar to the D423A-HSP60 cell system, many mitochondrial matrix and inner membrane proteins were decreased in amounts in HSP60 knockout zebrafish larvae, indicating that this effect of HSP60 deficiency is widespread across different tissues **(Figure 4J)**. The effect on mitochondrial matrix protein levels in fibroblast cells from patients carrying disease-associated HSP60 missense variations was very limited in P1 and undetectable in P2. This may be explained by the fact that the dominant-negative and knockout mutations severely shut down HSP60 function in D423A-HSP60 cells and zebrafish larvae, whereas the missense variations in the patients retain residual HSP60 function ^14,15^ and therefore have a much less pronounced effect on the mitochondrial proteome **(Figures 3A and 3B)**.

OXPHOS deficiency has been shown to trigger the mitochondrial integrated stress response (ISR^mt^) in a series of cellular and mouse models ^29,32–37^. The ISR^mt^ progresses in three stages, starting with the initial activation of FGF21-driven responses ^37^. Accordingly, in our D423A-HSP60 cell system, we observed upregulation of numerous genes that mark the different stages of the ISR^mt^, including *FGF21*, *ATF5*, *GDF15*, *MTHFD2*, *ASNS* from the first stage; *ATF3*, *PHGDH*, *PSAT1*, *SHMT2*, *CTH* and *TRIB3* from the second stage; and *DDIT3* from the third stage **(Figure 2H)**. Upregulation of zebrafish orthologs of ISR^mt^ genes was observed in our Hsp60 knockout zebrafish larvae through upregulation of *psat1*, *atf5b*, *phgdh*, *trib3* and *atf3* transcripts **(Figure 4I)**.

Consistent with these findings, metabolic analysis in both the D423A-HSP60 cell system and zebrafish Hsp60 knockout larvae showed elevated levels of serine and threonine **(Figures 5E and 5F)**. These are markers for alterations of the one-carbon cycle that includes the ISR^mt^ gene products PHGDH and PSAT1. Increased levels of serine and threonine have previously been reported for a mitochondrial respiratory chain dysfunction model ^29^. These metabolites generated inside mitochondria are important for the synthesis of building blocks of nucleic acids and for robust cell proliferation. Our metabolite analysis of induced D423A cells at 72h also showed decreased aspartate, which is the substrate for asparagine synthetase (ASNS) that converts it to asparagine. ASNS was upregulated both at protein and transcript levels in our cell model **(Figure 2G)**. Asparagine couples mitochondrial respiration to the activity of the ISR^mt^ transcription factor ATF4 ^38^ and supplementation of aspartate can rescue cell proliferation in electron transport deficient cells ^39^. Importantly, in HSP60 deficiency, the transcriptional upregulation of genes of the ISR^mt^ is inefficient because several of the encoded proteins (MTHFD2, SHMT2) fail to increase due to HSP60 chaperone deficiency. This prevents the ISR^mt^ from working properly.

The most striking observation in our study was the downregulation of the mevalonate and steroid pathways in both our D423A-HSP60 cell system and the zebrafish Hsp60 knockout model. Cholesterol is sensed by a system in the ER membrane that includes sterol regulatory element binding proteins (SREBPs) and insulin induced gene 1 (INSIG1) protein, which regulate promoters of genes involved in cholesterol biosynthesis, the LDL receptor (LDLR), ^40^, and fatty acid synthesis ^41^. This system senses the levels of both cholesterol and unsaturated fatty acids ^42^; low levels up-regulate and high levels down-regulate the expression of cholesterol synthesis genes. Our finding of downregulation of a large number of genes encoding enzymes of the mevalonate/cholesterol synthesis pathway and the LDL receptor at both transcript and protein levels in both the cellular and zebrafish models suggests that accumulation of cholesterol and/or fatty acids may occur. Indeed, oil red staining revealed highly elevated lipid levels in the yolk/liver region of zebrafish Hsp60 knockout larvae. Similar staining has been reported for knockdown of the alpha subunit of electron transfer flavoprotein (ETF) ^43^ that takes up electrons from more than ten flavoenzymes involved in fatty acid oxidation and amino acid degradation in the mitochondria ^44^. Accumulation of fatty acids due to reduced levels of fatty acid oxidation, together with impaired amino acid catabolism and mitochondrial pyruvate metabolism due to reduced levels of subunits of these will also lead to a shortage of acetyl-CoA, a key node in metabolism and an essential metabolite for epigenetic regulation ^45,46^.

These effects provide insight into the mechanisms underlying the hypomyelination phenotype in patients with HSP60 deficiency. Cholesterol is an abundant component of myelin sheaths - in humans, 25% of total body cholesterol is found in the brain, most of it as a component of myelin ^47^. In mammals, cholesterol used for building the myelin sheaths in the developing brain of the fetus is supplied by the mother from umbilical cord blood ^48^, and in zebrafish embryos cholesterol is provided by the yolk ^49^. Thus, there is no shortage of cholesterol during early brain development. However, once the blood-brain barrier is established, cholesterol for myelination must be synthesized locally in the CNS by astrocytes^19^. In humans, this occurs around birth, and in zebrafish, the blood-brain barrier begins to form at around 3 dpf and is fully functional at 10 dpf ^50^. Cholesterol availability from the yolk is the likely reason why we did not observe severe myelination phenotypes in zebrafish embryos by 8 dpf.

The synthesis of cholesterol and other lipids requires acetyl-CoA as a building block. Acetyl-CoA for the mevalonate/cholesterol pathway is derived from mitochondrial citrate, which is transported to the cytosol where it is converted back to acetyl-CoA by ATP-citrate lyase (ACLY) ^51^. Thus, once the blood-brain barrier is formed, the CNS in HSP60 deficient organisms face a double challenge for myelination: lack of the building block acetyl-CoA due to impaired catabolism of fatty acids, amino acids, and pyruvate in mitochondria and a down-regulation of cholesterol synthesis **(Figure 7)**. Hypomyelination would be expected to be more pronounced in severe HSP60 deficiency and less pronounced in milder HSP60 deficiency. The earliest brain MRI of a patient with the severe lethal form at the age of 5 months showed abnormal myelination, suggesting that myelination was impaired already at an early age ^15^. Clinical investigations of P1, with a mild presentation of HSP60 deficiency, revealed later onset and slow progression of disease phenotypes ^14^. The first MRIs of this patient, taken at age 4 years, showed delayed myelination of the cerebral and cerebellar white matter. A second MRI, taken at age 10 years, showed mild progression of his white matter disease **(Figure S7)**. This suggest that a mild HSP60 deficiency would lead to impaired remyelination/myelin maintenance after birth,

In summary, our extensive studies have shown that the levels of many mitochondrial matrix and inner membrane proteins are decreased in amounts when HSP60 is deficient. Our compendium of these data can be used to pinpoint mitochondrial proteins, enzyme complexes, and functions that are affected by dysregulation of HSP60 expression as seen e.g. in tumor cells, heart and skeletal muscle cells in cardiovascular disease, and diabetes ^52–55^.

The secondary effects of multiple deficiencies in the mitochondrial matrix and inner membrane proteins mount the ISR^mt^. However, in HSP60 deficiency, execution of the ISR^mt^ is hampered because the levels of the matrix proteins encoded by the up-regulated ISR^mt^ transcripts are not accordingly increasing. This results in an inappropriate - ‘limping’ - ISR^mt^.

As a major novel finding, we discovered a marked down-regulation, at both RNA and protein levels, of the cholesterol synthesis pathway upon acute shortage of HSP60. Hypomyelination and demyelination in the HSP60-related disorders appear thus to be due to impaired synthesis of cholesterol and related lipids in the brain. The roles of HSP60 activity in the wiring metabolism of lipids, amino acids, and other metabolites needs to be investigated in future studies. Novel treatments in HS60 deficiency and related hypo- or demyelinating diseases must address measures to counteract disadvantageous metabolic and regulatory responses affecting the cholesterol synthesis pathway.

Our study demonstrates the broad impact of HSP60 function on mitochondrial and cellular metabolism, maps the functions that are most strongly affected by HSP60 deficiency, and broadens the understanding of why tuning the expression and activity of this chaperone to cellular needs is of paramount importance for cellular homeostasis.

### Limitations of the Study

This study provides an important resource and understanding of HSP60 dysfunction in depth using three different models: (1) HEK cell model, (2) fibroblast cells from patients, (3) zebrafish animal model. On the other hand, some limitations must be considered. Firstly, one shortcoming is that the cell systems used do not include neuronal cells. In addition, neuronal cells only represent a small fraction of the larval body, which may mask nervous system-specific disease manifestations. However, the results derived from this study can be used as a guide to approach the tissue-specific pathogenesis characterizations in future studies. A common limitation of all animal models is that disease phenotypes may differ between species due to developmental and metabolic differences. An intrinsic problem of studying rare diseases using patient-derived fibroblasts is limited accessibility, as there are only very few known patients with disease-associated HSP60 mutations. However, the studies of rare diseases are particularly important for understanding and modeling the basic biology of complex human diseases.

## Material and Methods

### Cell Culture

HEK293 cells with tetracycline-inducible expression for wild-type (HSP60-WT) or the dominant-negative ATPase-deficient mutant (HSP60-D423A) were previously established using the Flp-In T-REx system (Invitrogen) ^56,57^. HSP60-WT and HSP60-D423A cells were sub-cultured in Dulbecco’s modified Eagle’s medium (DMEM, Sigma) supplemented with 10% fetal bovine serum, 0.29 mg/mL glutamine, 0.1 streptomycin mg/mL, 100 units penicillin/mL, 15 µg/mL blasticidin (Invitrogen) and 100 µg/ml hygromycin B (Invitrogen). Cells were cultured at 37°C, 85% humidity, and 5% CO_2_. Cells were routinely tested for Mycoplasma and grown in Mycoplasma-negative conditions. HSP60 expression (WT and D423A) was induced with 50 ng/mL tetracycline in culture media for the indicated induction periods. The medium was changed every other day and the day before measurements to maintain the tetracycline concentration constant in the culture medium.

### Cell Viability

HEK HSP60-WT and HSP60-D423A cells were cultured for 48, 72, and 96 hours and seeded at 1.2 million, 600.000, and 400.000 cells per T25 flask for the time points respectively. After trypsinization, cells were stained with acridine orange (AO; total cell staining) and DAPI (non-viable cell staining) and counted with a full-spectrum image cytometer, NucleoCounter NC-3000 (Chemometec, Denmark). The “Viability and Cell Count using NC-Slides™” Assay was run according to the manufacturer’s instructions. Cell viability was calculated by subtracting the non-viable cell number from the total cell number and represented as a percentage.

### Cell Proliferation

Cell proliferation rates were measured with label-free high-contrast brightfield microscopy in 6-well plates using Cytation 1 Cell Imaging Multi-Mode Reader (BioTek Instruments, Agilent) as described previously ^58^. Briefly, 125,000 cells were seeded per well, with or without tetracycline. Cell plates were incubated at room temperature for 30 minutes to promote uniform distribution of the attached cells, then incubated at 37°C with 5% CO_2_ until the next measurement. The cell medium was changed every other day after the measurement. Each sample was measured in three independent biological replicates, each set containing three wells of technical replicates in parallel. Once every 24 hours, two exact regions (9 auto-stitched images) from each well were imaged and analyzed. Cell proliferation rate was calculated by normalizing each cell count by the cell count at 24 hours post-seeding.

### Mitochondrial Mass Determination with MitoTracker™ Green

Mitochondrial mass was determined with MitoTracker™ Green FM (Invitrogen, USA, M7514). On the day of the analysis, cells were washed with PBS and incubated with 100 nM MitoTracker Green in HBSS (HankśBalanced Salt Solution) for 30 minutes at 37°C. Stained cells were trypsinized, collected into Eppendorf tubes, and stained with RedDot2 (Biotium, USA) (1:1000) to detect dead cells. The measurement was performed immediately using the NucleoCounter NC-3000 (Chemometec, Denmark) image cytometer. The total cell number was determined using the darkfield channel. Mean fluorescence intensity was calculated by the NucleoView software, dividing total fluorescence intensity by the total number of live cells.

### Mitochondrial Membrane Potential

Mitochondrial Membrane Potential Assay was run on an NC-3000 image cytometer (Chemometec), according to the manufacturer’s instructions. Briefly, 1×10^6^ cells/mL were stained with JC-1 (2.5 µg/mL) for 10 minutes at 37°C. Then, cells were washed with PBS and stained with DAPI solution (1 µg/mL) to detect dead cells and immediately analyzed. As a positive control of the assay, HEK HSP60-WT cells were treated with a mitochondrial uncoupler, Carbonyl Cyanide Chlorophenyl (CCCP; 50 µM), which is known to dissipate mitochondrial membrane potential.

### Mitochondrial Superoxide Detection with MitoSOX™ Red

Mitochondrial superoxide levels were determined with MitoSOX™ Red (Invitrogen, USA, M36008) using the image cytometer NC-3000, as described previously ^59^. Briefly, 1.2×10^6^ cells were seeded in 6-well plates 24 hours before the measurement with 5 µM MitoSOX™ Red for 30 minutes at 37°C. Un-induced HSP60-WT cells treated with 150 µM antimycin were used as positive control. Mean fluorescence intensity was calculated using the NucleoView software, by dividing total fluorescence intensity to the total number of living cells.

### Mitochondrial Bioenergetics

Mitochondrial bioenergetics was performed using Seahorse XFe96 (Seahorse, Agilent, USA) extracellular flux analyzer to measure Oxygen consumption rate (OCR) and extracellular acidification rate (ECAR) simultaneously. An hour before the measurement, the culture medium was replaced with the assay medium and incubated in a non-CO_2_ environment for 1 hour at 37°C. After the completion of OCR and ECAR measurements, each well was added Hoechst nuclear staining (2.7 µM/final well, Invitrogen, H3570) for 30 minutes in the dark at 37°C. Wells were imaged by Cytation1 using Cell Imaging software (Agilent) and OCR and ECAR values were normalized to the Hoechst-stained cell counts. The normalization unit was set to 1,000 cells.

For the ATP rate assay, we seeded 12,500 (uninduced), 15,000 (48h), and 17,500 (72h) cells/well. 48 hours before the assay, cells were seeded to 100 µg/mL Cell-Tak (Corning, #354241) pre-coated 96-well XF Cell Culture Microplates (Agilent; #101085-004). The assay medium was prepared by supplementing Seahorse XF DMEM medium (Agilent, #103575-100) with 10mM glucose (Sigma, G8769), 1 mM Sodium Pyruvate (Gibco, 11360-039), and 2 mM Glutamine (Sigma, G7513). ATP-synthetase inhibitor Oligomycin (1.5 µM final/well) and complex I and complex III inhibitors Rotenone and Antimycin A, respectively (0.5 µM final/well, Agilent, #103592-100), were injected sequentially. ATP production rate, mitoATP production rate, and glycoATP production rate were calculated by and exported from Wave Software (Agilent) according to the manufacturer’s instructions.

For the Mito Stress Test assay (Agilent, #103015-100), 25,000 cells from each cell line and condition were seeded to 100 µg/mL Cell-Tak pre-coated 96-well XF Cell Culture Microplates 24 hours before the assay. On the measurement day, the assay medium was prepared by supplementing XF Base medium, (Agilent, #103193-100; adjusted to pH 7.4) with 10 mM glucose (Sigma, G8769), 1 mM Sodium Pyruvate (Gibco, 11360-039), and 2 mM Glutamine (Sigma, G7513). Two group of sequential drug injections were used: (1) FCCP (Carbonyl cyanide 4-(trifluoromethoxy) phenylhydrazone; 0.25 µM final/well), and Rotenone/Antimycin A (0.5 µM final/well); (2) Oligomycin (1 µM final/well), and Rotenone/Antimycin A (0.5 µM final/well). The parameters obtained were calculated using the average of the three cycles measured according to the manufacturer recommendations.

### Bioluminescence ADP/ATP Ratio Determination

ADP/ATP ratio was determined in cell seeded into poly-D lysine (5 µg/mL, Sigma, P6407) pre-coated 96-Well Flat-Bottom White Microplates (Nunc, #136101). Between 2,000-4,000 cells were seeded to have 10.000 cells/well on the day of measurement. The culture medium was changed two hours before the measurement. ADP/ATP Ratio Assay Kit (Sigma, MAK135) was used according to the manufacturer’s instructions. Briefly, 90 µl of ATP reagent was added to each well and ATP-dependent bioluminescence (RLU_1_) was measured after one minute as relative light units (RLU) by Synergy H1 plate reader (BioTek, Agilent). Two wells without cells were used as a blank (RLU_0_). After 10 minutes of incubation in the dark, luminescence was re-measured (RLU_2_) to serve as a background for ADP measurement. Then, 5 µl of ADP reagent was added and ADP-dependent luminescence (RLU_3_) was measured. Measurements were performed in triplicates. ADP/ATP ratio was calculated as: (RLU_3_-RLU_2_)/ (RLU_1_-RLU_0_).

### Zebrafish Husbandry

To study early developmental Hsp60 defects in an animal model, we generated an *hspd1* knockout zebrafish line using CRISPR/Cas9 (Clustered Regularly Interspaced Short Palindromic Repeats (CRISPR)/CRISPR-associated protein 9). Adult zebrafish lines were housed and maintained at 28°C with 14/10 hours light/dark cycle. Embryos were maintained at 28.5°C in E3 buffer (5 mm NaCl, 0.17 mm KCl, 0.33 mm CaCl2, 0.33 mm MgSO4, 10−5% methylene blue, 2 mM HEPES, pH 7.4) with daily replacements. Wild-type zebrafish were in AB background. Crosses with Tg(*mbpa*:Dendra2) and Tg(*mbpa*:Dendra2-CaaX) transgenic lines were used for monitoring oligodendrocytes or myelin sheaths, respectively ^60^.

### sgRNA Selection

CHOPCHOP web tool (version 2; http://chopchop.cbu.uib.no/) was used to identify the potential target sites in the zebrafish *hspd1* gene and design the single guide RNAs (sgRNAs) used to select the target ^61^. The web tool lists the possible sgRNA sequences according to the protospacer adjacent motif (PAM) sequences located in a gene sequence, provides information on the potential efficiency, and reports the off-targets of the sgRNAs. We selected and tested a total number of 10 pairs of sgRNAs targeting exon 2 of *hspd1* to generate the *hspd1* knockout zebrafish line **(Table S1)**.

### sgRNA *In Vitro* Transcription

For the transcription of sgRNAs, pDR274 plasmid (Addgene; #42250) was linearised with the BsaI restriction enzyme (New England Biolabs; #R0535). The linearised plasmid was run on agarose gel and purified with QIAquick Gel Extraction Kit (Qiagen; #28704). sgRNA oligonucleotides (TAG Copenhagen AS, **Table S1**) were ligated to the linearised pDR274 plasmid, and cloned plasmids were transformed into XL-1 Blue cells (Agilent). Transformed cells were plated on Luria–Bertani (LB) medium agar plates supplemented with 50 µg/ml Kanamycin. Single colonies were selected and inoculated in LB medium, and DNA was extracted for Sanger sequencing to confirm the sgRNAs sequences. After confirming the sequences, the plasmids were linearised by the DraI restriction enzyme (New England Biolabs; R0129), a dual cutter for the plasmid. The band containing the T7 promoter and sgRNA template was cut from the agarose gel and purified with QIAquick Gel Extraction Kit (Qiagen; #28704). This linear DNA template containing the T7 promoter was used to transcribe sgRNAs with the MEGAshortscript™ T7 Transcription Kit (Invitrogen; AM1354). The synthesized gRNAs were precipitated with ethanol and dissolved in nuclease-free water.

### Injections of Zebrafish Embryos

Microinjections were performed by injecting 2 nL of sgRNA and Cas9 mRNA mixture into the yolk of 1-cell stage wild-type embryos from AB background. The efficiencies of the 10 pairs of sgRNAs on genomic DNA sequences from 1-day post-injection embryos were tested using the TIDE software (https://tide-calculator.nki.nl/) ^62^. To produce the F0 generation, the two most efficient sgRNA pairs were co-injected (50 pg of each sgRNA pair and 250 pg Cas9 mRNA) to 1-cell stage wild-type zebrafish embryos. Genomic DNA of the F0 generation’s somatic cells was obtained by skin-swabbing. The genetically altered adult F0 founder fish were outcrossed to wild-type zebrafish individually. The F1 generation was genotyped at 1-day post-fertilization (dpf) for the insertion/deletion (in/del) mutations in the germ cells. The F0 founder fish with in/dels resulting in frameshift mutations was outcrossed to generate F1 generation. Later, the adult F1 generation was mated to the wild-type background, resulting in F2 generation that constitute the heterozygous *hspd1* knockout line. Further generations were generated by outcrossing the *hspd1*^+/-^ knockout line **(Figure S3A and S3B)**.

### Brightfield Imaging

The F4 generation of the *hspd1*^+/-^ knockout line was in-crossed, and brightfield imaging of larvae was performed at 3 dpf, 5 dpf, and 7 dpf with Olympus SZX16 (Olympus) equipped with Axiocam 305 colour (Zeiss), set at 2x zoom. Larvae were anesthetized in 225 mg/mL buffered tricaine (pH 7.2), imaged in glass-bottom dishes (MatTek; #P35G-1.0-20-C) and placed on to a thin layer of 3% methylcellulose in E3. Imaged larvae were carefully placed back into an individual well of a p24-well plate and maintained with daily E3 replacements until the next measurement point. At the end of 7 dpf imaging, larvae were euthanized and genotyped. Body length, swim-bladder area, and eye area were measured as shown **(Figure S3C)** using ImageJ.

### Oil Red O staining

The zebrafish larvae resulting from hspd1^+/-^ in-cross were euthanized in tricaine at 8 dpf and washed twice with PBS before overnight fixation with 4% PFA at 4°C. Fixed larvae were incubated in 60% isopropanol for one hour at room temperature. 0.3% Oil Red O (ORO) in 60% isopropanol was freshly prepared and filtered through a 0.2 µM syringe filter. ORO solution was added to the fixed larvae and incubated for 3 hours at room temperature with a gentle shake. Larvae were washed with 60% isopropanol twice and PBS with 0.1% Tween was added. Larvae were transferred sequentially to 25% glycerol, 50% glycerol, and finally to 75% glycerol in PBS with 0.1% Tween, and directly imaged with Olympus SZX16 (Olympus) dissection microscope equipped with Axiocam 305 color (Zeiss) at 6.3x zoom.

### Fluorescent Imaging

The F4 generation *hspd1*^+/-^ knockout line was crossed to two transgenic zebrafish lines fluorescently labeled for oligodendrocytes, Tg(*mbpa*:Dendra2); and myelin sheaths, Tg(*mbpa*:Dendra2-CaaX). *hspd1*^+/-^ Tg(*mbpa*:Dendra2) line or *hspd1*^+/-^ Tg(*mbpa*:Dendra2-CaaX) line were in-crossed. The resulting larvae were anesthetized in 225 mg/mL buffered tricaine (pH 7.2), imaged in glass-bottom dishes (MatTek; #P35G-1.0-20-C), embedded in 1% low-melting agarose and kept in individual wells of a p24-well plate in-between measurements. The larvae were blindly imaged with an Observer.Z1 fluorescent microscope (Zeiss, Germany) equipped with AxioCam HRm camera, ApoTome.2 slider, and 10x objective, image stacks were acquired in ZEN 2.3 software (Zeiss) in the FITC channel, the maximum intensity projections (MIP) from images of 2 µm optical sections were generated with ImageJ (version 1.53c) software, and MIPs were stitched manually. Dendra2-positive mature oligodendrocytes were blindly quantified from the caudal to the yolk sac extension. Only viable larvae were imaged, quantified, and genotyped after the analysis.

### TMT Labeling and Mass Spectrometry-Based Proteomics Sample Preparation

For protein isolation from HEK cells, HSP60-D423A cells were seeded in T75 flasks with the following cell numbers: uninduced (3.6 million), 48h (3.6 million), and 72h (2.4 million) induced. The human dermal fibroblast cells, control and patient fibroblasts were cultured in T75 flasks (250,000 cells/flask) for 48 hours. On the day of the collection, cells were washed with warm PBS once, 3 mL ice-cold PBS was added to the flasks, and cells were detached with a cell scraper (VWR, #734-2602) and collected into a pre-chilled falcon tube in 3 repeated steps. Samples were centrifuged at 400 g for 3 minutes at 4°C, the supernatant was removed, and cell pellets were stored at -80°C until lysis.

The F4 generation of the zebrafish *hspd1*^+/-^ knockout line was in-crossed and obtained embryos were maintained with daily E3 replacements until 5 dpf. The 5 dpf larvae were anesthetized with 150 mg/L tricaine in groups of four at a time. The tip of the caudal fin of the anesthetized larvae was then removed using a hypodermic needle. The larvae body was immediately snap-frozen in an empty 96-well plate (Biorad; #12001925) on dry ice and stored at -80°C until sample preparation. The tail was transferred to another 96-well plate with 100% methanol-filled wells for genotyping. According to the obtained genotyping results, bodies of *hspd1*^+/+^ and *hspd1*^-/-^ larvae were pooled in groups of 5 larvae, and this process was repeated for 5 mating sets, resulting in 10 samples in total **(Figure S4A)**.

HEK cell model pellets, zebrafish animal model larvae and fibroblast cell pellets were lysed in 1%SDS in 100 mM triethylammonium bicarbonate (TEAB). Zebrafish larvae were additionally homogenized by a motorized pellet pestle (Kimble; #Z359971). All samples were further lysed by sonication with a microtip sonicator (Branson Sonifier 250; Branson Ultrasonics) at 30% duty cycle for three cycles of five pulses with a minute of ice incubation between cycles. Sonicated lysates were centrifuged at 16,000 g for 10 minutes, and supernatants were transferred to a new tube. Protein concentrations were determined with the BCA protein assay (Thermo Scientific; #10678484).

### Tandem Mass Tag Labeling

HEK cells and zebrafish samples were tagged with TMT10plex™ Isobaric Label Reagent Set (Thermo Scientific, #90111), and fibroblast cells with TMTpro™16plex Isobaric Labels (Thermo Scientific, #A44520). From each sample, 80 µg total protein was in-solution trypsin digested and TMT labeled according to the manufacturer’s instructions. Following the TMT labeling, all samples within each study were pooled. After a strong cation exchange (SCX) purification (Phenomenex, #8B-S010-EAK), peptides were loaded to an Immobiline™ DryStrip Gel (GE Healthcare, #11534985) isoelectric focusing (IEF) separation was performed. The IEF gels were then cut into 10 equal pieces, purified in PepClean C18 spin columns (Thermo Scientific, #11824141) according to the manufacturer’s instructions, and then vacuum centrifuged until dryness and stored at -20°C until nanoLC-MS/MS analyses. For the HEK cells, 8.89 µg of un-labeled total protein from all samples were pooled and labeled as a separate sample and used to normalize TMT ratios for each experimental sample.

### NanoLC-MS/MS and Proteomics Database Searches

Nano-liquid chromatography tandem-mass spectrometry (nanoLC-MS/MS) was performed on an EASY nanoLC-1200 coupled to a Q-Exactive™ HF-X Quadrupole-Orbitrap™ Mass Spectrometer (Thermo Scientific) as previously described ^63^. Briefly, MS was operated in positive mode using pre- and analytical columns: acclaim PepMap 100, 75 µm × 2 cm (Thermo Scientific) and EASY-Spray PepMap RSLC C18, 2µm, 100 Å, 75 µM × 25 cm (Thermo Scientific), respectively, to trap and separate peptides with a 170-minute gradient in 5-40 % acetonitrile, 0.1 % formic acid. Higher-energy collisional dissociation (HCD) was used for peptide fragmentation, and the normalized collision energy (NCE) was 35. Full scan (MS1) and fragment scan resolutions were set at 60,000 and 45,000, respectively. Automatic gain control (AGC) for MS1 and MS2 were set at 1×10^6^ with a scan range between 392-1,500 m/z and at 1×105 with a fixed mass of 110 m/z, respectively. Up to 12 of the most intense peaks of the full scan were fragmented with data-dependent analysis (DDA). Unassigned and +1 charge states were excluded from fragmentation, and the dynamic exclusion duration was 15 seconds. Each fraction was LC-MS/MS analyzed twice, where peptides identified from >9 scans in the first analysis were excluded from fragmentation in the second analysis.

### Proteomics Database Search

All LC-MS/MS results were merged and submitted for the database search for identification and quantification of proteins using Proteome Discoverer 3.0 (Thermo Scientific). For the HEK and fibroblast cell models, 20,401 reviewed Homo sapiens Uniprot sequences were used as reference proteome (Swiss-Prot; downloaded on 02.12.2022) using the Sequest algorithm. For the zebrafish animal model, 46,691 *Danio rerio* Uniprot sequences were used as reference proteome (UP000000437; downloaded on 02.12.2022). Precursor mass tolerance was 10 ppm, and fragment mass tolerance was 20 mmu. The maximum number of allowed missed cleavages was two. Oxidation of methionine was set as dynamic modification and static modifications were carbamidomethylation of cysteines and TMT-plex-labels on lysine and peptide N-terminus. The co-isolation threshold was set at 35%, and the identification false discovery rate was 0.01 at both peptide and protein levels. Proteins with more than 2 quantitative peptide scans and at least one unique peptide were considered as quantified and included in the further proteomics data analysis. Criteria for the differentially expressed proteins were set at p<0.05 and |FC|>1.2 for protein abundance.

### RNA Sequencing Sample Preparation

HEK cell pellets were obtained as described in proteomics sample preparation. Differently, after the last centrifugation step, PBS was removed, 250 µL of TRIzol (Invitrogen, #15596-018) was added, and the sample snap-frozen on dry ice, and stored at -80°C for later use for RNA sequencing.

Larvae of the zebrafish animal model were obtained from in-crosses of the F4 generation of the *hspd1*^+/-^ knockout line and collected as described in the proteomics section. Larvae were covered 25 µL TRIzol (Invitrogen; #15596-018) and stored at -80°C until RNA extraction. RNA was extracted from pools of 8 whole-body *hspd1*^+/+^ and *hspd1*^-/-^ larvae at 5 dpf. This process was repeated for 3 mating sets, resulting in 6 samples in total. Pooled samples were homogenized with a motorized pellet pestle (Kimble; #Z359971).

RNA isolation was performed using TRIzol, including DNase treatment with DNA-free Kit (Ambion, AM1906) according to the manufacturer’s instructions. Total RNA concentrations were determined with Synergy H1 plate reader equipped with Take3 Plate (BioTek, Agilent). Quality control analysis was performed with an automated 4200 TapeStation System (Agilent, G2991AA), and RNA integrity number (RIN) values were determined by RNA ScreenTape Assay for 4200 TapeStation System according to the manufacturer’s instructions. RIN values were above 9.3 and 8.3 for all samples of the cell and zebrafish models, respectively. RNA sequencing service was performed by BGI Copenhagen by a non-stranded and polyA-selected RNA library preparation and a consequent PE100 sequencing on DNBSEQ. Data was obtained from BGI in fastq file format. Initial quality control of the fastq files was conducted using FastQC (Babraham Bioinformatics). Adaptor removal and trimming of low-quality ends were performed using Trim Galore with default settings (Babraham Bioinformatics). Gene expression was quantified using Salmon against decoy-aware reference transcriptomes (GRCh38 and *Danio rerio*, GRCz11) ^64^. Transcript abundances were calculated and summarized at the gene level with tximeta ^65^. Differential expression analysis was performed using DESeq2 ^66^, applying a false discovery rate (FDR) threshold of less than 0.05.

### Mass Spectrometry-Based Metabolomics Sample Preparation

HSP60-D423A cell line cells were seeded to T75 flasks, uninduced (3.6 million cells), 48h (3.6 million cells), and 72h (2.4 million cells). Cell culture flasks were placed onto a 37°C warm plate, medium was removed, cells were washed with warm PBS once, PBS was discarded, 4 mL of 80% Methanol (-20°C, VWR, #85800290) was directly added to cells with the flasks on ice. Cells were detached with a cell scraper (VWR, #734-2602), collected into a pre-chilled falcon tube in 3 repeated steps, vortexed for 5 seconds, and incubated on ice for 10 minutes. The samples were sonicated in a water bath for 10 seconds with 3 repeated cycles with 1 minute of ice incubations in between. For the zebrafish metabolomics study, larvae at 5 dpf were obtained from the in-cross of the F5 generation of the hspd1^+/-^ knockout line and collected as above-described in the proteomics section. Pools of 8 larvae were homogenized in ice-cold 80% methanol using a motorized Kimble pellet pestle and sonicated for 10 seconds in a water bath.

The metabolite extracts were centrifuged at 12.000 g for 10 minutes at 4°C. The supernatants were evaporated to dryness by a flow of nitrogen gas and then re-dissolved in water:methanol (97:3). TCA metabolites citric acid, alpha-ketoglutaric acid, malic acid, succinic acid, glutamic acid, and lactate were measured by LC-MS/MS as described previously ^67^. Briefly, the samples were prepared by mixing 25 µL re-dissolved supernatant with 80 µL of stable isotope-labeled internal standard solution (D4-citric acid, D4-succinic acid, D4-α-ketoglutaric acid, D3-malic acid, and D3-lactic acid) dissolved in water with 0.3 % formic acid. For the analysis of amino acids, 10 µL supernatant was added 40 µL solution of stable isotope-labeled internal standards (phenylalanine-D5, tyrosine-D4, valine-D8, methionine-D3, lysine-13C6-15N2, and arginine 13C6-15N4; Cambridge Isotope Laboratories) and 180 µL solvent (water with 0.2% heptafluorobutyric acid). Reference material amino acids (certified amino acid mix, Sigma Aldrich) were used to prepare calibrator samples in the concentration range 0.2 to 25 µM by addition of internal standard and solvent as specified above. Three µl of the mixture was injected into an ultra-performance liquid chromatography system (Waters UPLC) coupled to a mass spectrometer (Waters Xevo TQ-S). The column was an HSS T3 (Waters Acquity UPLC 100 x 2.1 mm, 1.8 µm, 100 Å) maintained at 40 °C. The initial flow rate was 0.6 mL/min of 99% mobile phase A (water with 0.1% heptafluorobutyric acid) and 1% B (acetonitrile) that was maintained for 1 min, followed by a gradient to 20% B (1.0-4.6 min) to 80% B (4.6-5.5 min) and then back to 1% B (5.5-6.0 min) with equilibration for 2 min, giving a total run time of 8 min. The mass spectrometer used electrospray ionization in the positive mode with capillary voltage 3.0 kV, source temperature 150 °C, desolvation temperature 600 °C, and 800 L/h desolvation gas flow (nitrogen). The amino acids were detected using multiple reaction monitoring with the following m/z transitions, glycine (76→30.1), aspartate (134→74), alanine (90.1→44.2), threonine (120.1→74.2), glutamine (147→84.2), serine (106→60.2), asparagine (133.1→87.2), valine (118.1→72.2), phenylalanine (166.1→120.2), leucine (132.1→86.2), lysine (147.1→84.2), arginine (175.1→70.2), methionine (150→104.1), tyrosine (182.1→136.2), phenylalanine-D5 (171→125), tyrosine-D4 (186→146), valine-D8 (126→80), methionine-D3 (153→107), lysine-13C6-15N2(155→90), and arginine 13C6-15N4(185.1→75.2). Calibration curves were constructed by linear regression of the peak area ratio (amino acid/internal standard) versus the nominal concentrations of the calibrator samples with a weighting factor of 1/X. Selected stable isotope analogs were employed for quantification of their corresponding amino acids and other amino acids according to retention time proximity. Data for TCA metabolites and amino acids were normalized against the total ion intensity of each cell supernatant sample as obtained from an untargeted metabolomics LC-MS analysis of the sample by a qTOF instrument (Bruker Q-TOF maXis Impact) combined with XCMS-based peak analyses as described previously ^68^.

### HSP60 Peptide Levels by Mass Spectrometry-Based Proteomics

Lysates from the proteomics study were used to quantify HSP60-WT and HSP60-D423A protein amounts. Briefly, 30 µg total protein was loaded on AnykD™ Criterion™ TGX™ Precast Midi Protein Gel (Biorad, #5671125) and 60 kDa region was excised from the Coomassie stained gel. Gel pieces were destained and in-gel trypsin digested (Promega, V5280). The peptide mixture was purified on PepClean C18 spin columns (Thermo Scientific, #11824141) and analyzed with nano Liquid-Chromatography coupled to a Q-Exactive mass spectrometer (Thermo Fisher Scientific). Detected HSP60 peptides were identified and quantitated using Proteome Discoverer version 2.3 and MaxQuant software. HSP60-D423A protein expression levels were estimated by calculating the ratio of the endogenously expressed HSP60-WT peptide (VTDALNATR) to all the other quantified HSP60 peptides.

### Transcript Expression Quantification by RNA-sequencing

*HSPD1* and *hspd1* transcript expression levels were quantified by quasi-mapping as implemented in the Salmon software program ^64^ in HEK cells and zebrafish, respectively. A salmon index was built based on WT and Mutant *HSPD1* transcripts (Ensembl, GRCh38.p13) for HEK cells and WT transcripts (ENSDART00000078596.6 and ENSDART00000127938.3) and KO transcripts (Ensembl, GRCz11) for zebrafish, using default parameters. Next, these transcripts were quantified in all samples (paired-end reads, processed as described in the RNAseq section above). The “salmon quant” command was evoked with default parameters, and the WT to mutant ratio was calculated.

### Western blotting

Embryos were lyzed with CelLytic M lysis buffer (Sigma; C2978) supplemented with cOmplete™, Mini Protease Inhibitor Cocktail (Roche; 11836153001) according to manufacturer’s instructions. Protein concentrations were determined by Bradford reagent (Biorad; #5000205) measurements performed in flat-bottom 96-well plates in triplicates with Synergy H1(BioTek, Agilent) plate reader at 595 nm wavelength absorbance. Bovine Serum Albumin (BSA) standard curve was used to calculate protein concentrations of the samples. 30 µg of total protein was loaded to AnykD™ Criterion™ TGX Stain-Free™ Protein Gels (BIORAD; #5678125). Transfer of the proteins from gel to the Immun-Blot® Low Fluorescence PVDF membrane with low fluorescent transfer buffer (BIORAD; #1704275) was performed in Trans-Blot Turbo Transfer System (BIORAD) at 10V for 30 minutes. Once the transfer was completed, the blot was visualized under UV for the total protein measurement and later used to normalize the detected antibody signal. Antibody-treated blots were developed with Enhanced Chemiluminescence Plus (ECL; Pierce; 32132), and Chemiluminescence or Cy2 fluorescence signals were recorded.

### Bioinformatics and Enrichment Analysis

R was used to merge and edit data tables and to add information from Human MitoCarta3.0 and MitoPathways3.0 (https://www.broadinstitute.org/mitocarta/mitocarta30-inventory-mammalian-mitochondrial-proteins-and-pathways). ID conversion and alignment for human proteins, genes and transcripts was done based on HGNC Biomart (https://biomart.genenames.org/martform). For Zebrafish proteomics to RNASeq comparisons, ID alignment tables of ZFIN Marker associations to Ensembl IDs, NCBI Gene data, and UniProt protein data were downloaded from zfin.org. Zfish/human orthology comparisons are based on Zfin Human and Zebrafish Orthology from zfin.org. Vulcano plots were generated in R using ggplot. Split violin plots were generated using ggplot added the introdataviz package (https://psyteachr.github.io/introdataviz/advanced-plots.html).

For the zebrafish animal model study, orthologous human genes were identified by Biomart and used at ENSDAR and ENSEMBL IDs levels. For the analysis of mitochondrial proteins, ENSEMBL IDs of the human MitoCarta3.0 ^5^ mitochondrial inventory was used to detect and match the respective orthologous human genes of zebrafish ENSDAR IDs ^69^.

Enrichment analysis of the differentially expressed proteins or transcripts (proteins p-value <0.05 and |log2FC|>0.26, and transcripts p-adj.value <0.05 and |log2FC|>1) was performed on the Enrichr and FishEnrichr, for cell and zebrafish models respectively ^27,28^. Analyses were performed at the gene name level for proteomics and transcriptomics. The enriched terms are shown as a negative log10 of p-values from the Fisher exact test.

### Statistics

The mean, standard deviation, standard error of the mean, and student’s t-test were calculated in Excel or Prism 9.0 (GraphPad Software). The indicated statistical tests were performed using GraphPad Prism version 9.1.2 for Windows. The p-value of p<0.05 was considered significant in statistical calculations.

### Ethics Statement

The Danish Animal Experiments Inspectorate approved the generation of the *hspd1* knockout zebrafish animal model (Project nr: 2018-15-0201-01451). Zebrafish were handled according to Danish legislation at all times.

The use of human dermal fibroblasts study conformed to the Declaration of Helsinki. The Central Denmark Region Committees on Health Research Ethics determined proteomic analysis of the patient fibroblasts not to be a Human Health Research Study, thus Institutional review board approval was not required. Informed consent for the use of the MRI and clinical data was obtained previous to their use.

## Resource availability

### Lead contact

Further information and requests for resources and reagents should be directed to and will be fulfilled by the lead contact Peter Bross.

### Materials availability

This study did not generate new unique reagents.

### Data and code availability

Code: No original code was generated in this study.

Any additional information required to reanalyze the data reported in this paper is available from the lead contact upon request.

All data will be shared upon request. The human data will be shared anonymized.

## Acknowledgments

We thank Helle Highland-Nygaard and Margrethe Kjeldsen (Research Unit for Molecular Medicine, Aarhus University and Aarhus University Hospital, Denmark) for their technical assistance, and Tianran Zhou and Xiaqing Yu for help with handling and extraction of transcriptomics data. The RNA sequencing analysis was performed by BGI Copenhagen, Denmark.

This work was supported by a PhD fellowship from the Graduate School of Health, Aarhus University, Denmark (C.C.). We also thank the Aarhus County Research Initiative, Lundbeck Foundation (R315-2018-2521 to C.C.), Max and Magda Nørgaard Foundation (WZ632-0044/20-2000 to C.C.), the Eva & Henry Frænkels Mindefond (P.B.), and the European Molecular Biology Organization (Short-Term Fellowship #8638 to C.C.) for support.

## Author contributions

C.C., P.F.G., and P.B. conceived the study and designed most of the experiments. C.C. performed most of the experiments. C.C. and P.F.G. performed and analyzed the mitochondrial function, cellular characterization, and respirometry of the D423A-HSP60 cellular system. C.C. and J.C. performed and analyzed cell proliferation measurements. C.C., K.K.S., and L.S.L. designed, performed, and analyzed the zebrafish experiments. J.J., J.P., J.C., Y.L., and C.O. provided key resources and critical scientific advice. BFM and EØ provided human fibroblast samples, and contributed clinical information, C.C. and J.H. performed and analyzed the metabolomics experiments. C.C., J.J., and P.B. performed and analyzed the transcriptomics studies. C.C., J.P., and P.B. performed and analyzed the proteomics studies. C.C., P.F.G., and P.B. analyzed the data, wrote and revised the manuscript. All authors contributed to data interpretation, read, edited and approved the final version of the manuscript.

## Declaration of Interests

The authors declare no competing interests.

## Supplemental data

**Table S1:**
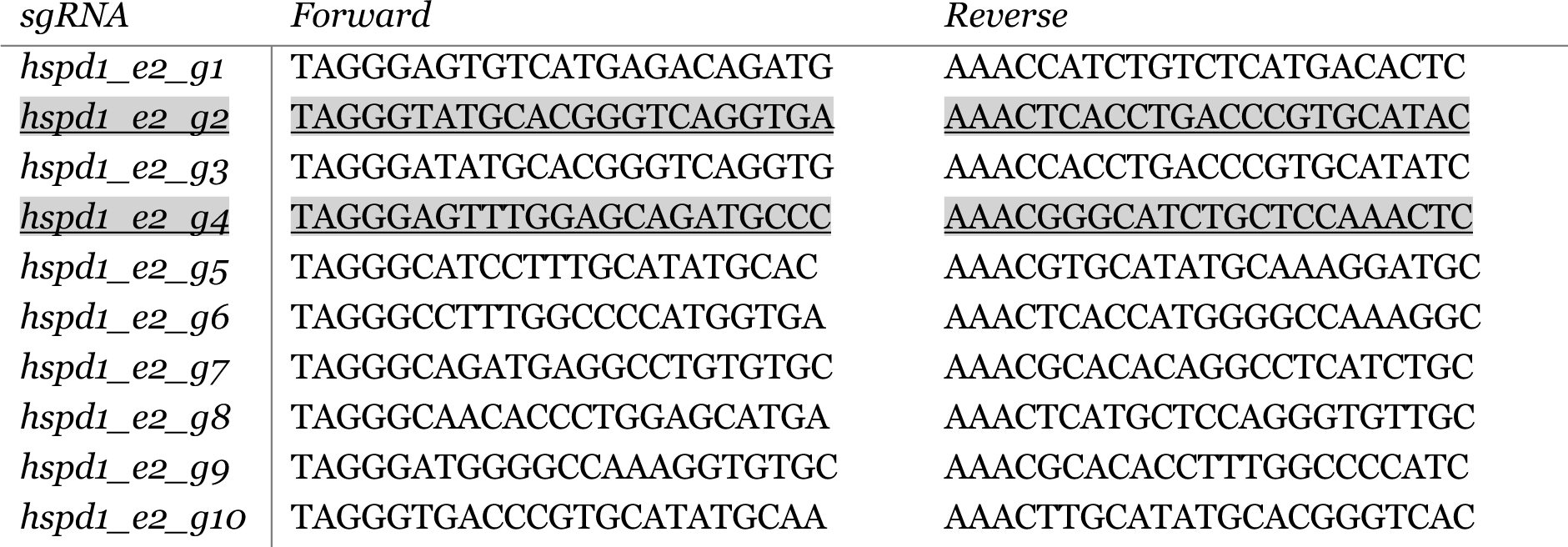
sgRNA sequences, most efficient sgRNA pair highlighted in gray. hspd1_e2_g2 had 18.6% efficiency for 200 pg/embryo and hspd1_e2_g4 had 71.5% efficiency for 200 pg/embryo.

**Figure S1:**
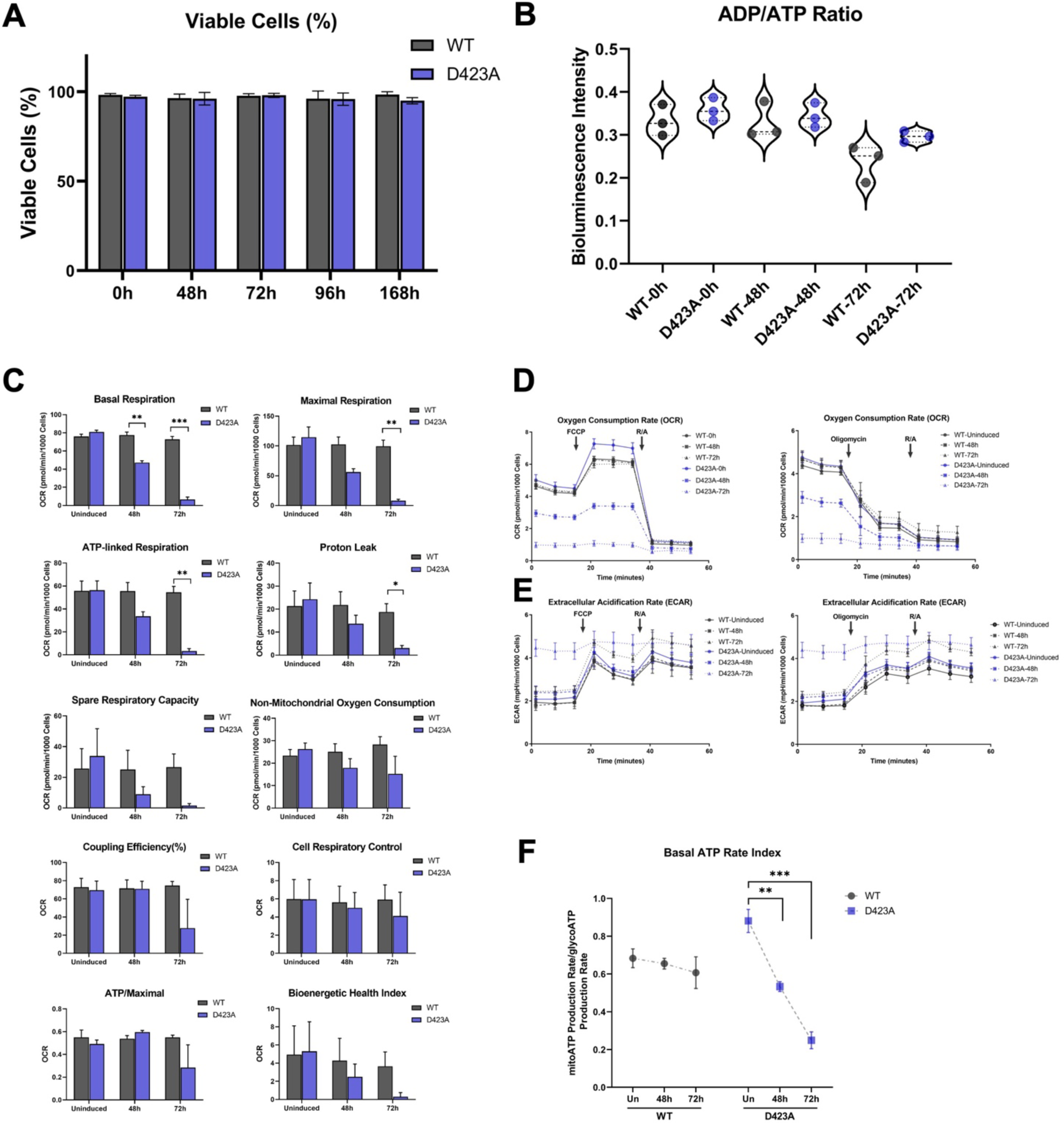
**(A)** Viable cell percentage is represented as the percentage of live cells to total cells. Measured at 0h, 48h, 72h, 96h, and 168h induced WT and D423A cells (mean ±SD, n≥3) **(B)** ADP/ATP ratio was determined by a bioluminescence-based assay at 0h, 48h, and 72h induced WT and D423A cells (n = 3). **(C)** Graphs showing basal respiration, maximal respiration, ATP-linked respiration, proton leak, spare respiratory capacity, non-mitochondrial oxygen consumption, coupling efficiency in percentage, cell respiratory control, ATP/Maximal (ATP-linked respiration to maximal ratio), and Bioenergetic health index were calculated following the Mito Stress Test Assay according to the manufacturer’s instructions. **(D)** Representative OCR measurement trace graphs of 0h, 48h, and 72h induced WT (shades of grey) and D423A (shades of blue) cells at baseline and after mitochondrial inhibitor injections (FCCP, Oligo, R/A). **(E)** Representative ECAR measurement trace graphs of 0h, 48h, and 72h induced WT (shades of grey) and D423A (shades of blue) cells at baseline and after mitochondrial inhibitor injections (FCCP, Oligo, R/A). **(F)** Mitochondrial to glycolytic ATP production rates ratio was measured in the presence of the mitochondrial inhibitors (Oligo, R/A) and calculated from OCR and PER (mean ±SD, n=3, multiple unpaired t-tests with Holm-Sidak correction). Abbreviations: OCR, oxygen consumption rate; ECAR, extracellular acidification rate FCCP, carbonyl cyanide 4-(trifluoromethoxy)phenylhydrazone; Oligo, Oligomycin; R/A, Rotenone and Antimycin A; ADP, adenosine diphosphate; ATP, adenosine triphosphate.

**Figure S2:**
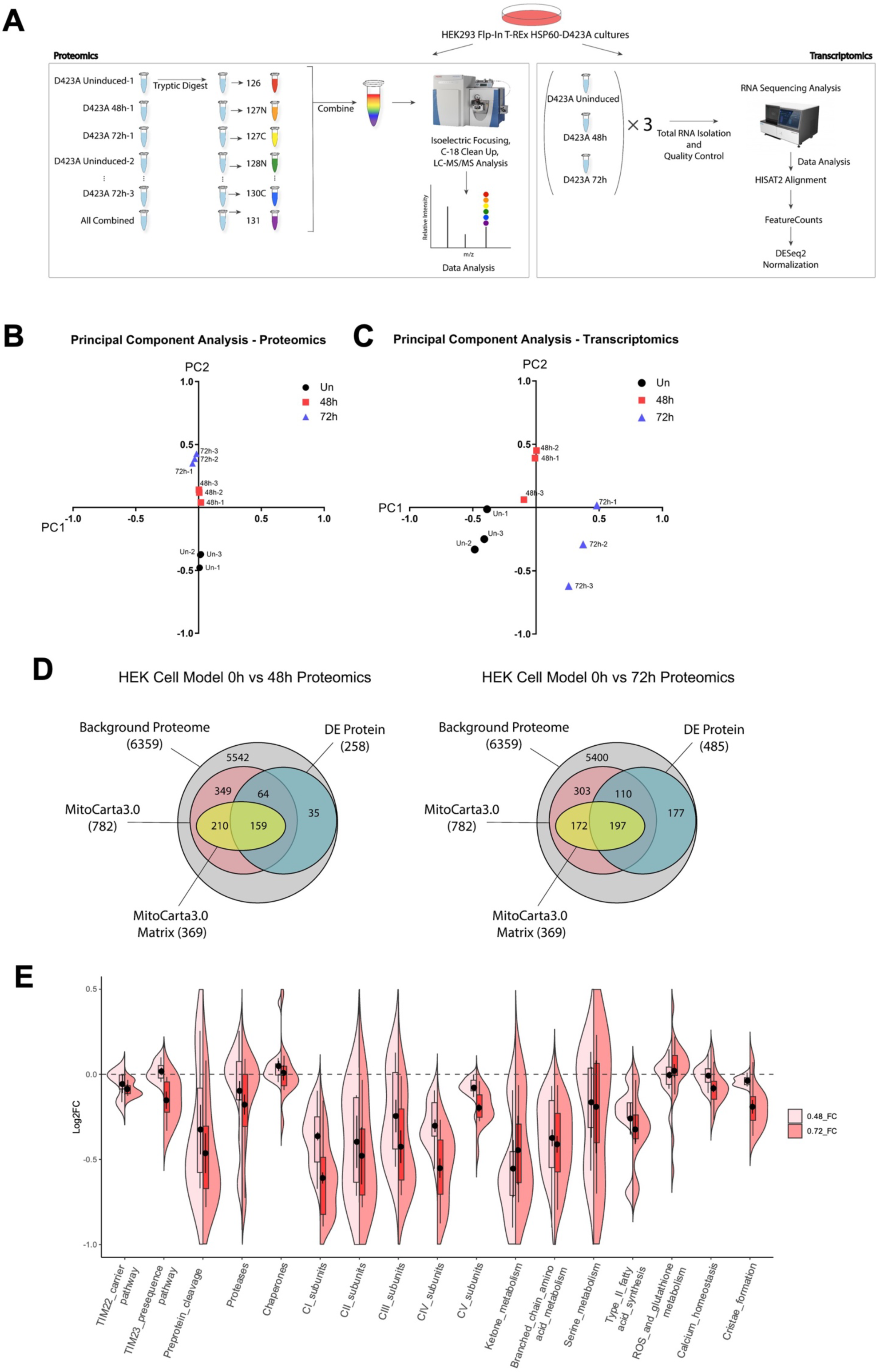
**(A)** Schematic of experimental design summarizing the proteomics and transcriptomics of D423A cells; 0h, 48h, and 72h induced. **(B-C)** Principal component analysis (PCA) of proteomics and transcriptomics analysis of D423A cells; 0h, 48h, and 72h induced. **(D)** Venn diagrams representing the number of quantified mitochondrial proteins based on MitoCarta3.0 and indicating differentially expressed proteins (Proteomics: p-value <0.05 and |log2FC|>0.26). Uninduced D423A cells were compared to 48h (left) or 72h induced D423A cells (right). The pairing of quantified proteins was performed through Ensembl gene IDs. **(E)** Violin plots distinguishing proteins from a selection of mitochondrial MitoCarta pathways of induced D423A cells at 0h in comparison to 48h (light red) or 72h induced D423A cells (dark red).

**Figure S3:**
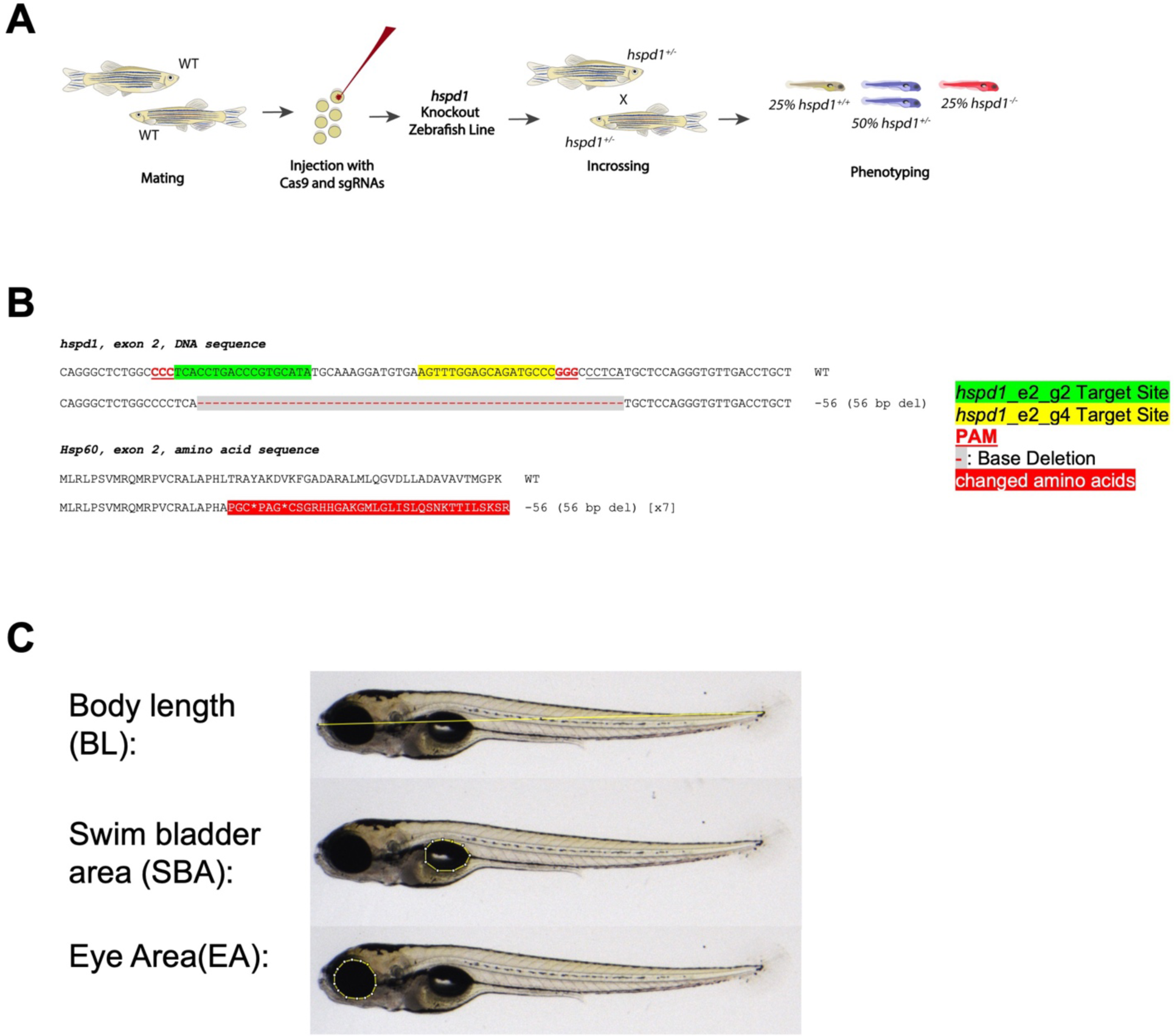
**(A)** Schematic showing the main steps of CRISPR/Cas9-mediated generation of *hspd1* knockout. Briefly, ten sgRNA pairs were screened for efficiency by injecting wild-type embryos with sgRNAs and Cas9 mRNA. The two most efficient sgRNA pairs were co-injected and resulting mosaic embryos were raised to adulthood (F0 generation). F0 adults were individually screened for germline transmission, and germline-confirmed F0 adults were individually outcrossed to generate the F1 generation. Once the hspd1 knockout is established in the F1 generation, the following generations were generated by outcrossing to WT adults, and resulting embryos were used for phenotyping. **(B)** Sequence summary for germline transmission of F0 generation to the F1 generation. F0 mosaic individuals were generated with 50 ng hspd1_e2_g2, hspd1_e2_g4 sgRNA, and Cas9 mRNA injection. F0 individuals were mated to the WT zebrafish line to determine the germline transmission sequence of the mosaic F0 individuals. The DNA sequence from exon 2 of the hspd1 gene and the respective amino acid sequences are shown. The crRNAs; hspd1_e2_g2 target site is shown in green, hspd1_e2_g4 sgRNA target site is indicated in yellow. The PAM sequences are underlined in red. Deleted basepairs are indicated with a red dash highlighted in grey. In the amino acid sequences, changed amino acids are highlighted in red. The resulting deletion is shown next to the sequence. The top line indicates the WT sequence. Deletions are indicated with a minus sign, and the insertion (in) or deletion (del) are specified in parenthesis. The indicated 56 bp deleted allele constitutes the hspd1 knockout zebrafish line. **(C)** Quantification of body length, head-trunk angle, size of the inflated swim bladder, and eye size were determined as shown in the figure using ImageJ.

**Figure S4:**
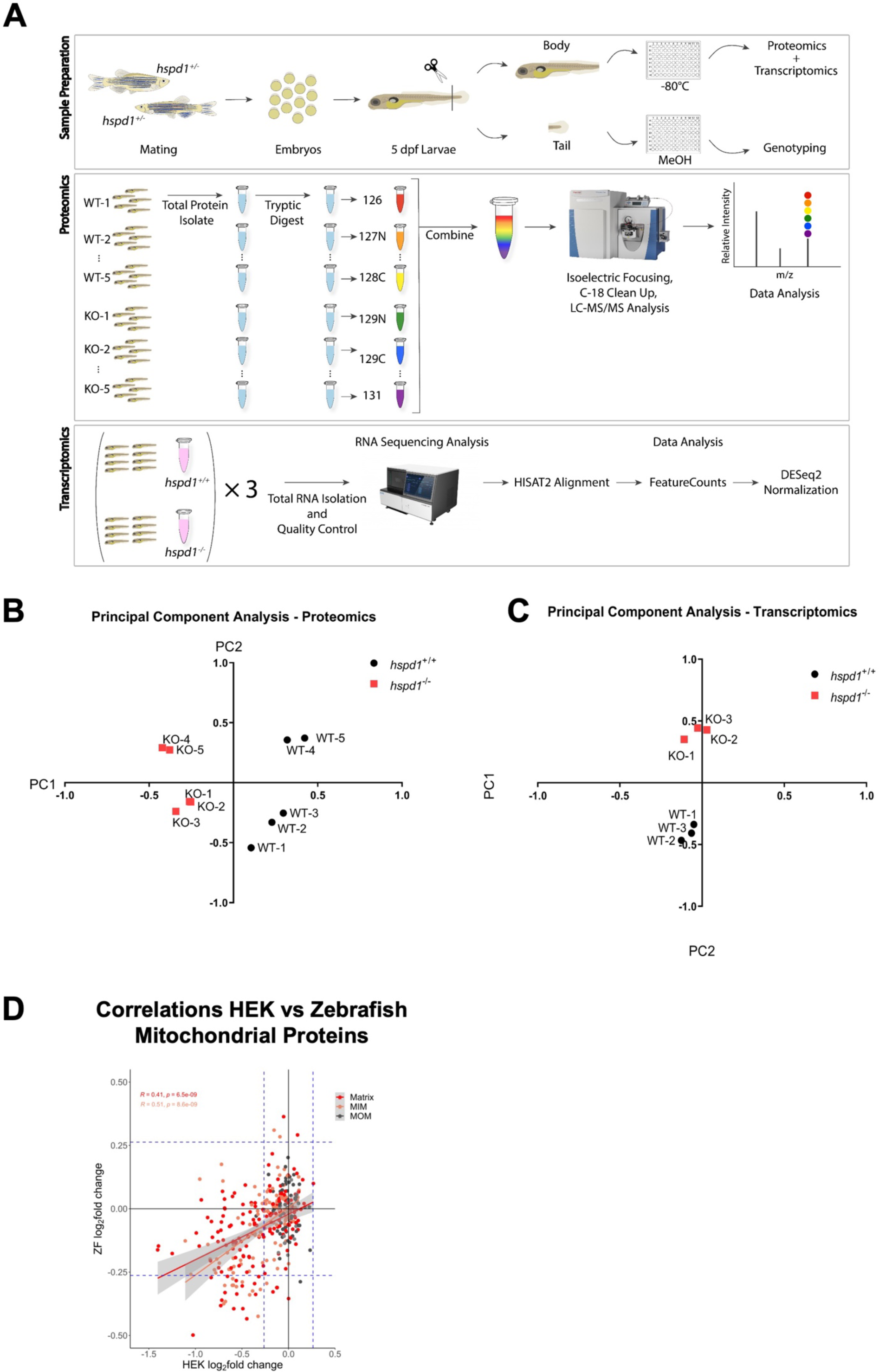
**(A)** Experimental workflow for the proteomic and transcriptomic characterization of *hspd1* knockout in zebrafish larvae. Adult *hspd1^+/-^* fish were mated, and embryos were raised to 5 dpf. Larval tails were cut and used for genotyping, and bodies were stored at -80°C and used for proteomics or transcriptomics studies. For proteomics, groups of 5 larvae were pooled per sample in 5 replicates (10 samples). For transcriptomics, groups of 8 larvae were pooled per sample in 3 replicates (6 samples). **(B-C)** Principal component analysis of proteomics (**B**) and transcriptomics (**C**) indicating clustering of *hspd1^+/+^* (black, WT) and *hspd1^-/-^* (red, KO) replicate samples. **(D)** Correlation of log2 fold changes of mitochondrial proteins in the D423A-HSP60 cells (log2 fc 0h versus 72h induction; horizontal axis) versus the Zebrafish HSP60 knockout larvae (log2 fc wt versus ko; vertical axis). Pearson *R* and *p* values for the correlation for points associated with the subcompartments are given and correlation lines for matrix and inner membrane proteins are shown. Abbreviations: See figure 2 and 4.

**Figure S5:**
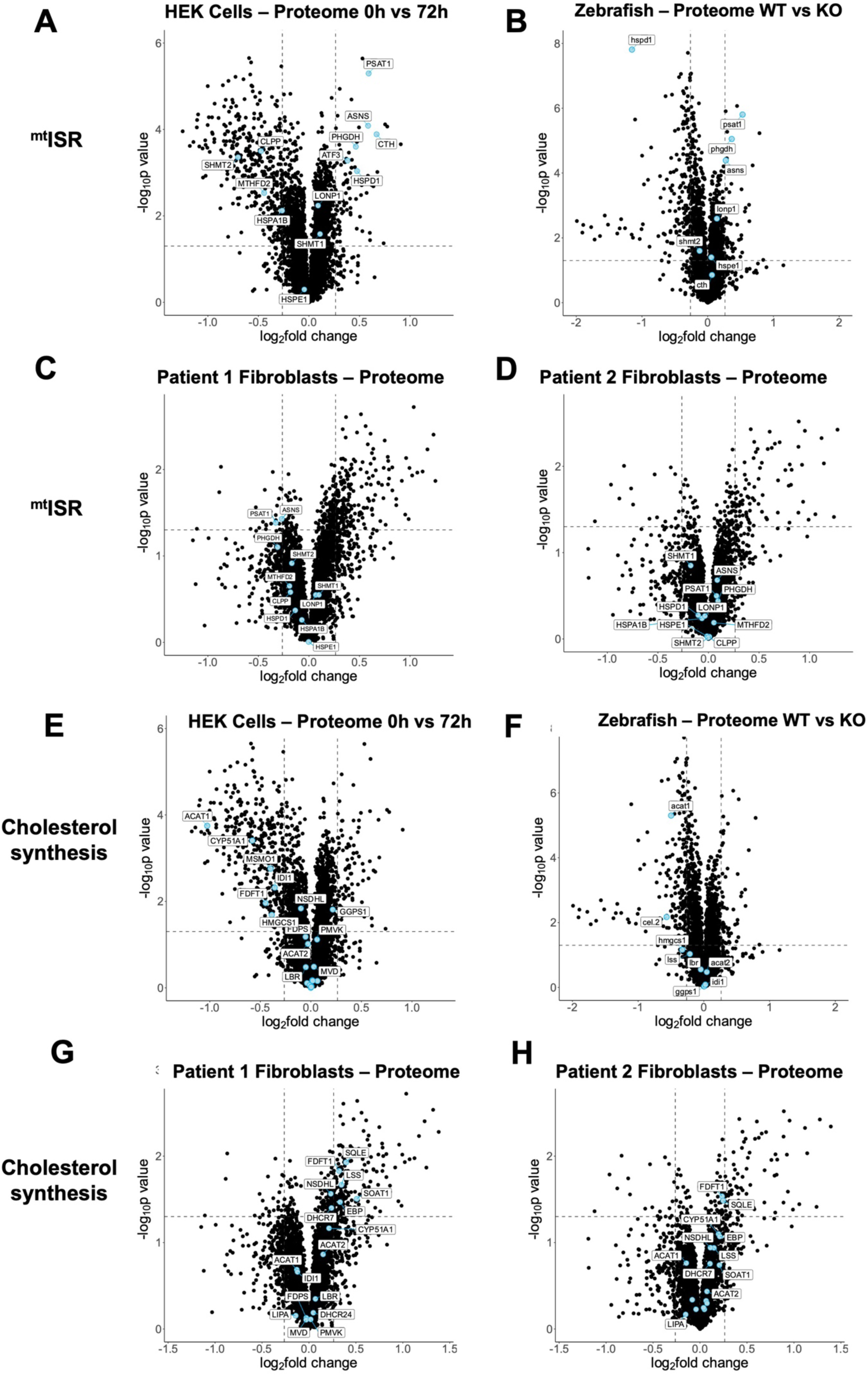
Protein changes of the ISR^mt^ and cholesterol synthesis. **(A-D):** Volcano plots showing the regulation of mitochondrial integrated stress response (ISR^mt^) proteins in D423A-HSP60 cells induced for 72h **(A),** in 5 dpf zebrafish *hspd1^-/-^* larvae **(B),** in fibroblasts from patient 1 **(C),** and fibroblasts from patient 2 **(D)**. **(E-H)** Volcano plots showing the regulation of proteins involved in cholesterol synthesis pathways in D423A-HSP60 cells induced for 72h **(E),** in 5 dpf zebrafish *hspd1^-/-^* larvae **(F)**, in fibroblast cells from patient 1 **(G)** and patient 2 **(H).**

**Figure S6:**
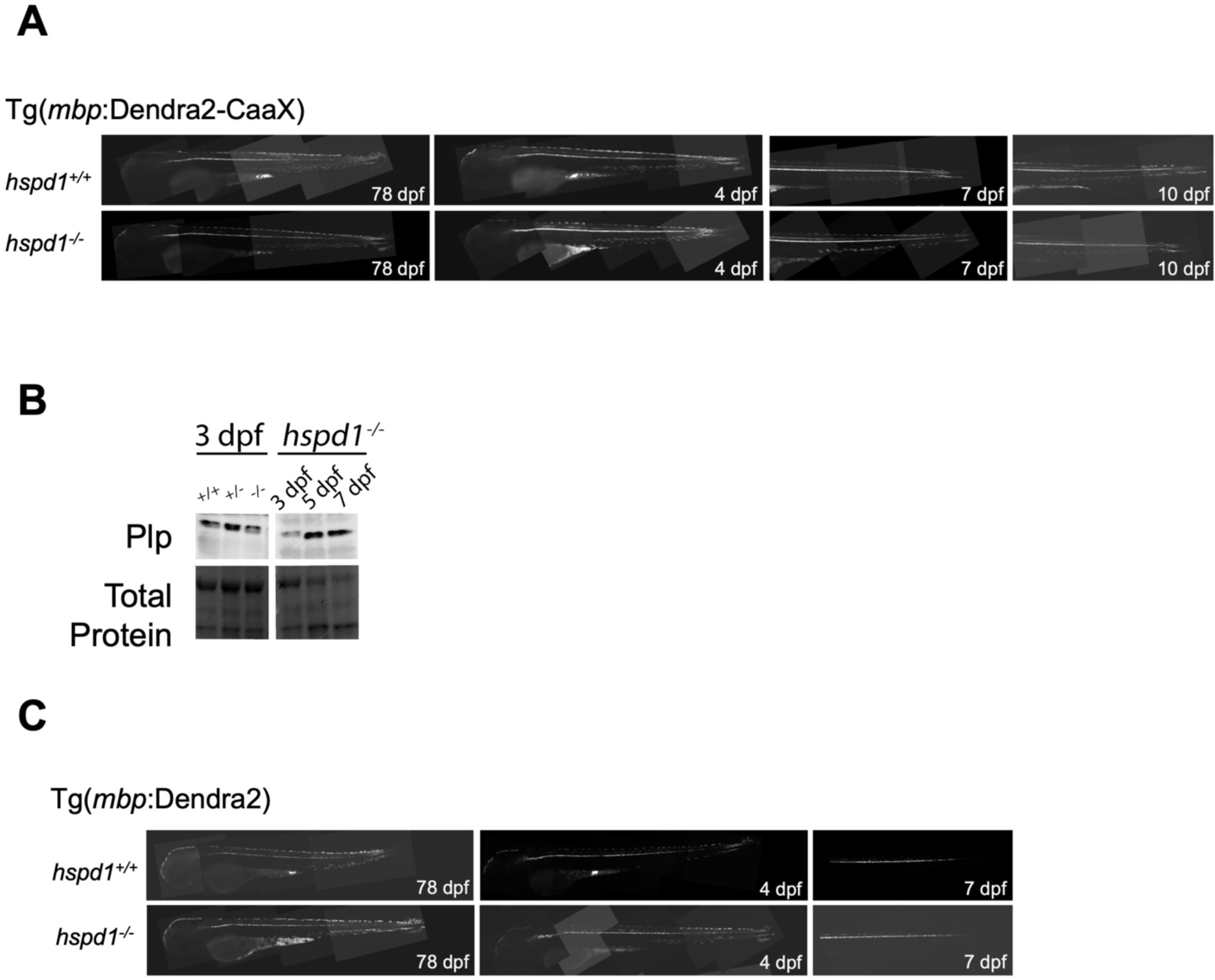
**(A)** Myelination of *hspd1^+/+^*, *hspd1^+/-^*, and *hspd1^-/-^* larvae was studied at 4 dpf, 7 dpf, and 10 dpf in Tg(*mbpa*:Dendra2-CaaX) background. **(B)** Plp protein levels were assessed in the absence of Hsp60 in zebrafish larvae by western blot protein quantification of *hspd1^+/+^*, *hspd1^+/-^*, and *hspd1^-/-^* larvae at 3 dpf and *hspd1^-/-^* larvae at 3, 5, and 7 dpf. **(C)** Mature oligodendrocytes of *hspd1^+/+^*, *hspd1^+/-^*, and *hspd1^-/-^* larvae were studied at 78 hpf, 4 dpf and 7 dpf in Tg(mbpa:Dendra2) background.

**Figure S7:**
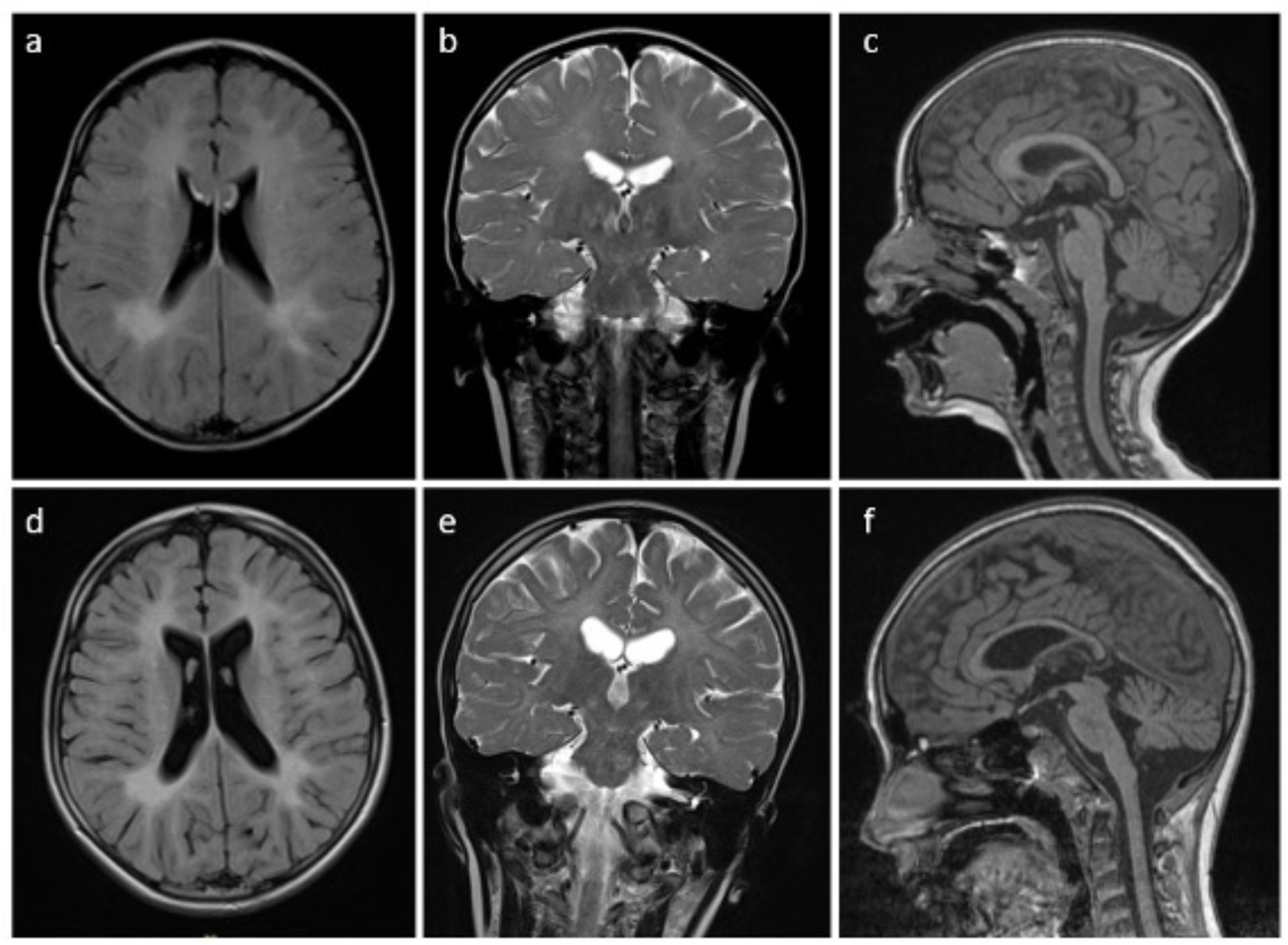
MR images of Patient 1 at age 4 (**a, b, c**) and 10 years (**d, e, f**) show abnormal signal intensity in the cerebral and cerebellar white matter, extending to involve the subcortical U fibers, with hyperintensity on FLAIR (**a, d**) and T2 (**b, e**), and normal signal on T1(c,f), in keeping with hypomyelination. Follow-up images show progression over time with an increase in the confluent peritrigonal abnormal signal, interval thinning of the corpus callosum, mild enlargement of the lateral and third ventricles, and prominence of the cerebral sulci in keeping with atrophy.

## Notes

### Competing Interest Statement

The authors have declared no competing interest.

## References

1. Monzel, A.S., Enriquez, J.A., and Picard, M. (2023). Multifaceted mitochondria: moving mitochondrial science beyond function and dysfunction. Nat Metab 5, 546–562. 10.1038/s42255-023-00783-1.

2. Raichle, M.E., and Gusnard, D.A. (2002). Appraising the brain’s energy budget. Proc Natl Acad Sci U S A 99, 10237–10239. 10.1073/pnas.172399499.

3. Johri, A., and Beal, M.F. (2012). Mitochondrial dysfunction in neurodegenerative diseases. J. Pharmacol. Exp. Ther. 342, 619–630. 10.1124/jpet.112.192138.

4. Traxler, L., Lagerwall, J., Eichhorner, S., Stefanoni, D., D’Alessandro, A., and Mertens, J. (2021). Metabolism navigates neural cell fate in development, aging and neurodegeneration. Dis Model Mech 14. 10.1242/dmm.048993.

5. Rath, S., Sharma, R., Gupta, R., Ast, T., Chan, C., Durham, T.J., Goodman, R.P., Grabarek, Z., Haas, M.E., Hung, W.H.W., et al. (2021). MitoCarta3.0: an updated mitochondrial proteome now with sub-organelle localization and pathway annotations. Nucleic Acids Res 49, D1541–D1547. 10.1093/nar/gkaa1011.

6. Kim, Y.E., Hipp, M.S., Bracher, A., Hayer-Hartl, M., and Hartl, F.U. (2013). Molecular chaperone functions in protein folding and proteostasis. Annu Rev Biochem 82, 323–355. 10.1146/annurev-biochem-060208-092442.

7. Saibil, H. (2013). Chaperone machines for protein folding, unfolding and disaggregation. Nat Rev Mol Cell Biol 14, 630–642. 10.1038/nrm3658.

8. Bie, A.S., Comert, C., Korner, R., Corydon, T.J., Palmfeldt, J., Hipp, M.S., Hartl, F.U., and Bross, P. (2020). An inventory of interactors of the human HSP60/HSP10 chaperonin in the mitochondrial matrix space. Cell Stress Chaperones 25, 407–416. 10.1007/s12192-020-01080-6.

9. Christensen, J.H., Nielsen, M.N., Hansen, J., Fuchtbauer, A., Fuchtbauer, E.M., West, M., Corydon, T.J., Gregersen, N., and Bross, P. (2010). Inactivation of the hereditary spastic paraplegia-associated Hspd1 gene encoding the Hsp60 chaperone results in early embryonic lethality in mice. Cell Stress.Chaperones. 15, 851–863.

10. Pei, W., Tanaka, K., Huang, S.C., Xu, L., Liu, B., Sinclair, J., Idol, J., Varshney, G.K., Huang, H., Lin, S., et al. (2016). Extracellular HSP60 triggers tissue regeneration and wound healing by regulating inflammation and cell proliferation. NPJ Regenerative medicine 1. 10.1038/npjregenmed.2016.13.

11. Bross, P., and Fernandez-Guerra, P. (2016). Disease-Associated Mutations in the HSPD1 Gene Encoding the Large Subunit of the Mitochondrial HSP60/HSP10 Chaperonin Complex. Front Mol Biosci 3, 92–98. 10.3389/fmolb.2016.00049.

12. Kusk, M.S., Damgaard, B., Risom, L., Hansen, B., and Ostergaard, E. (2016). Hypomyelinating Leukodystrophy due to HSPD1 Mutations: A New Patient. Neuropediatrics 47, 332–335. 10.1055/s-0036-1584564.

13. Yamamoto, T., Yamamoto-Shimojima, K., Ueda, Y., Imai, K., Takahashi, Y., Imagawa, E., Miyake, N., and Matsumoto, N. (2018). Independent occurrence of de novo HSPD1 and HIP1 variants in brothers with different neurological disorders - leukodystrophy and autism. Hum Genome Var 5:18. 10.1038/s41439-018-0020-z.

14. Cömert, C., Brick, L., Ang, D., Palmfeldt, J., Meaney, B.F., Kozenko, M., Georgopoulos, C., Fernandez-Guerra, P., and Bross, P. (2020). A recurrent de novo HSPD1 variant is associated with hypomyelinating leukodystrophy. Cold Spring Harb Mol Case Stud 6. 10.1101/mcs.a004879.

15. Magen, D., Georgopoulos, C., Bross, P., Ang, D., Segev, Y., Goldsher, D., Nemirovski, A., Shahar, E., Ravid, S., Luder, A., et al. (2008). Mitochondrial hsp60 chaperonopathy causes an autosomal-recessive neurodegenerative disorder linked to brain hypomyelination and leukodystrophy. Am.J.Hum.Genet. 83, 30–42.

16. Wolf, N.I., ffrench-Constant, C., and van der Knaap, M.S. (2020). Hypomyelinating leukodystrophies — unravelling myelin biology. Nature Reviews Neurology. 10.1038/s41582-020-00432-1.

17. Escande-Beillard, N., Loh, A., Saleem, S.N., Kanata, K., Hashimoto, Y., Altunoglu, U., Metoska, A., Grandjean, J., Ng, F.M., Pomp, O., et al. (2020). Loss of PYCR2 Causes Neurodegeneration by Increasing Cerebral Glycine Levels via SHMT2. Neuron 107, 82–94 e86. 10.1016/j.neuron.2020.03.028.

18. Berghoff, S.A., Spieth, L., and Saher, G. (2022). Local cholesterol metabolism orchestrates remyelination. Trends Neurosci. 45, 272–283. 10.1016/j.tins.2022.01.001.

19. Saher, G., and Stumpf, S.K. (2015). Cholesterol in myelin biogenesis and hypomyelinating disorders. Biochim. Biophys. Acta 1851, 1083–1094. 10.1016/j.bbalip.2015.02.010.

20. Berger, E., Rath, E., Yuan, D., Waldschmitt, N., Khaloian, S., Allgauer, M., Staszewski, O., Lobner, E.M., Schottl, T., Giesbertz, P., et al. (2016). Mitochondrial function controls intestinal epithelial stemness and proliferation. Nature communications 7, 13171. 10.1038/ncomms13171.

21. Fan, F., Duan, Y., Yang, F., Trexler, C., Wang, H., Huang, L., Li, Y., Tang, H., Wang, G., Fang, X., et al. (2020). Deletion of heat shock protein 60 in adult mouse cardiomyocytes perturbs mitochondrial protein homeostasis and causes heart failure. Cell Death Differ 27, 587–600. 10.1038/s41418-019-0374-x.

22. Magnoni, R., Palmfeldt, J., Christensen, J.H., Sand, M., Maltecca, F., Corydon, T.J., West, M., Casari, G., and Bross, P. (2013). Late onset motoneuron disorder caused by mitochondrial Hsp60 chaperone deficiency in mice. Neurobiol Dis 54, 12–23. 10.1016/j.nbd.2013.02.012.

23. Bie, A.S., Palmfeldt, J., Hansen, J., Christensen, R., Gregersen, N., Corydon, T.J., and Bross, P. (2011). A cell model to study different degrees of Hsp60 deficiency in HEK293 cells. Cell Stress Chaperones. 16, 633–640.

24. Magnoni, R., Palmfeldt, J., Hansen, J., Christensen, J.H., Corydon, T.J., and Bross, P. (2014). The Hsp60 folding machinery is crucial for manganese superoxide dismutase folding and function. Free Radical Res 48, 168–179. 10.3109/10715762.2013.858147.

25. Corydon, T.J., Hansen, J., Bross, P., and Jensen, T.G. (2005). Down-regulation of Hsp60 expression by RNAi impairs folding of medium-chain acyl-CoA dehydogenase wild-type and disease-associated proteins. Mol.Genet.Metab 85, 260–270.

26. Karczewski, K.J., Francioli, L.C., Tiao, G., Cummings, B.B., Alföldi, J., Wang, Q., Collins, R.L., Laricchia, K.M., Ganna, A., Birnbaum, D.P., et al. (2020). The mutational constraint spectrum quantified from variation in 141,456 humans. Nature 581, 434–443. 10.1038/s41586-020-2308-7.

27. Kuleshov, M.V., Jones, M.R., Rouillard, A.D., Fernandez, N.F., Duan, Q., Wang, Z., Koplev, S., Jenkins, S.L., Jagodnik, K.M., Lachmann, A., et al. (2016). Enrichr: a comprehensive gene set enrichment analysis web server 2016 update. Nucleic Acids Res. 44, W90–97. 10.1093/nar/gkw377.

28. Xie, Z., Bailey, A., Kuleshov, M.V., Clarke, D.J.B., Evangelista, J.E., Jenkins, S.L., Lachmann, A., Wojciechowicz, M.L., Kropiwnicki, E., Jagodnik, K.M., et al. (2021). Gene Set Knowledge Discovery with Enrichr. Curr Protoc 1, e90. 10.1002/cpz1.90.

29. Bao, X.R., Ong, S.E., Goldberger, O., Peng, J., Sharma, R., Thompson, D.A., Vafai, S.B., Cox, A.G., Marutani, E., Ichinose, F., et al. (2016). Mitochondrial dysfunction remodels one-carbon metabolism in human cells. Elife 5. 10.7554/eLife.10575.

30. Chandel, N.S. (2021). Lipid Metabolism. Cold Spring Harb Perspect Biol 13. 10.1101/cshperspect.a040576.

31. Busch, J.D., Fielden, L.F., Pfanner, N., and Wiedemann, N. (2023). Mitochondrial protein transport: Versatility of translocases and mechanisms. Mol. Cell 83, 890–910. 10.1016/j.molcel.2023.02.020.

32. Forsstrom, S., Jackson, C.B., Carroll, C.J., Kuronen, M., Pirinen, E., Pradhan, S., Marmyleva, A., Auranen, M., Kleine, I.M., Khan, N.A., et al. (2019). Fibroblast Growth Factor 21 Drives Dynamics of Local and Systemic Stress Responses in Mitochondrial Myopathy with mtDNA Deletions. Cell Metab 30, 1040–1054 e1047. 10.1016/j.cmet.2019.08.019.

33. Kuhl, I., Miranda, M., Atanassov, I., Kuznetsova, I., Hinze, Y., Mourier, A., Filipovska, A., and Larsson, N.G. (2017). Transcriptomic and proteomic landscape of mitochondrial dysfunction reveals secondary coenzyme Q deficiency in mammals. Elife 6. 10.7554/eLife.30952.

34. Nikkanen, J., Forsstrom, S., Euro, L., Paetau, I., Kohnz, R.A., Wang, L., Chilov, D., Viinamaki, J., Roivainen, A., Marjamaki, P., et al. (2016). Mitochondrial DNA Replication Defects Disturb Cellular dNTP Pools and Remodel One-Carbon Metabolism. Cell metabolism 23, 635–648. 10.1016/j.cmet.2016.01.019.

35. Samluk, L., Urbanska, M., Kisielewska, K., Mohanraj, K., Kim, M.J., Machnicka, K., Liszewska, E., Jaworski, J., and Chacinska, A. (2019). Cytosolic translational responses differ under conditions of severe short-term and long-term mitochondrial stress. Mol Biol Cell 30, 1864–1877. 10.1091/mbc.E18-10-0628.

36. Ignatenko, O., Malinen, S., Rybas, S., Vihinen, H., Nikkanen, J., Kononov, A., Jokitalo, E.S., Ince-Dunn, G., and Suomalainen, A. (2023). Mitochondrial dysfunction compromises ciliary homeostasis in astrocytes. J Cell Biol 222. 10.1083/jcb.202203019.

37. Suomalainen, A., and Battersby, B.J. (2018). Mitochondrial diseases: the contribution of organelle stress responses to pathology. Nat. Rev. Mol. Cell Biol. 19, 77–92. 10.1038/nrm.2017.66.

38. Krall, A.S., Mullen, P.J., Surjono, F., Momcilovic, M., Schmid, E.W., Halbrook, C.J., Thambundit, A., Mittelman, S.D., Lyssiotis, C.A., Shackelford, D.B., et al. (2021). Asparagine couples mitochondrial respiration to ATF4 activity and tumor growth. Cell metabolism. 10.1016/j.cmet.2021.02.001.

39. Birsoy, K., Wang, T., Chen, W.W., Freinkman, E., Abu-Remaileh, M., and Sabatini, D.M. (2015). An Essential Role of the Mitochondrial Electron Transport Chain in Cell Proliferation Is to Enable Aspartate Synthesis. Cell 162, 540–551. 10.1016/j.cell.2015.07.016.

40. Shimano, H., and Sato, R. (2017). SREBP-regulated lipid metabolism: convergent physiology — divergent pathophysiology. Nature Reviews Endocrinology 13, 710–730. 10.1038/nrendo.2017.91.

41. Vergnes, L., Chin, R.G., De Aguiar Vallim, T., Fong, L.G., Osborne, T.F., Young, S.G., and Reue, K. (2016). SREBP-2-deficient and hypomorphic mice reveal roles for SREBP-2 in embryonic development and SREBP-1c expression. J. Lipid Res. 57, 410–421. 10.1194/jlr.m064022.

42. Madison, B.B. (2016). Srebp2: A master regulator of sterol and fatty acid synthesis. J. Lipid Res. 57, 333–335. 10.1194/jlr.C066712.

43. Kim, S.H., Scott, S.A., Bennett, M.J., Carson, R.P., Fessel, J., Brown, H.A., and Ess, K.C. (2013). Multi-organ abnormalities and mTORC1 activation in zebrafish model of multiple acyl-CoA dehydrogenase deficiency. PLoS Genet. 9, e1003563. 10.1371/journal.pgen.1003563.

44. Henriques, B.J., Katrine Jentoft Olsen, R., Gomes, C.M., and Bross, P. (2021). Electron transfer flavoprotein and its role in mitochondrial energy metabolism in health and disease. Gene, 145407. 10.1016/j.gene.2021.145407.

45. Shi, L., and Tu, B.P. (2015). Acetyl-CoA and the regulation of metabolism: mechanisms and consequences. Curr. Opin. Cell Biol. 33, 125–131. 10.1016/j.ceb.2015.02.003.

46. Martinez-Reyes, I., and Chandel, N.S. (2020). Mitochondrial TCA cycle metabolites control physiology and disease. Nature communications 11, 102. 10.1038/s41467-019-13668-3.

47. Bjorkhem, I., and Meaney, S. (2004). Brain cholesterol: long secret life behind a barrier. Arterioscler Thromb Vasc Biol 24, 806–815. 10.1161/01.ATV.0000120374.59826.1b.

48. Russell, D.W., Halford, R.W., Ramirez, D.M., Shah, R., and Kotti, T. (2009). Cholesterol 24-hydroxylase: an enzyme of cholesterol turnover in the brain. Annu. Rev. Biochem. 78, 1017–1040. 10.1146/annurev.biochem.78.072407.103859.

49. Fraher, D., Sanigorski, A., Mellett, N.A., Meikle, P.J., Sinclair, A.J., and Gibert, Y. (2016). Zebrafish Embryonic Lipidomic Analysis Reveals that the Yolk Cell Is Metabolically Active in Processing Lipid. Cell Rep 14, 1317–1329. 10.1016/j.celrep.2016.01.016.

50. Morris, E.K., Daignault-Mill, S., Stehbens, S.J., Genovesi, L.A., and Lagendijk, A.K. (2023). Addressing blood-brain-tumor-barrier heterogeneity in pediatric brain tumors with innovative preclinical models. Frontiers in oncology 13. 10.3389/fonc.2023.1101522.

51. Chandel, N.S. (2021). Mitochondria. Cold Spring Harb Perspect Biol 13. 10.1101/cshperspect.a040543.

52. Tang, Y., Zhou, Y., Fan, S., and Wen, Q. (2022). The Multiple Roles and Therapeutic Potential of HSP60 in Cancer. Biochem. Pharmacol., 115096. 10.1016/j.bcp.2022.115096.

53. Duan, Y., Tang, H., Mitchell-silbaugh, K., Fang, X., Han, Z., and Ouyang, K. (2020). Heat Shock Protein 60 in Cardiovascular Physiology and Diseases. Frontiers in molecular biosciences 7. 10.3389/fmolb.2020.00073.

54. Kleinridders, A., Lauritzen, H.P., Ussar, S., Christensen, J.H., Mori, M.A., Bross, P., and Kahn, C.R. (2013). Leptin regulation of Hsp60 impacts hypothalamic insulin signaling. J. Clin. Invest. 123, 4667–4680. 10.1172/JCI67615.

55. Hauffe, R., Rath, M., Schell, M., Ritter, K., Kappert, K., Deubel, S., Ott, C., Jähnert, M., Jonas, W., Schürmann, A., and Kleinridders, A. (2021). HSP60 reduction protects against diet-induced obesity by modulating energy metabolism in adipose tissue. Molecular Metabolism, 101276. 10.1016/j.molmet.2021.101276.

56. Bie, A.S., Palmfeldt, J., Hansen, J., Christensen, R., Gregersen, N., Corydon, T.J., and Bross, P. (2011). A cell model to study different degrees of Hsp60 deficiency in HEK293 cells. Cell Stress Chaperones 16, 633–640. 10.1007/s12192-011-0275-5.

57. Bross, P., Naundrup, S., Hansen, J., Nielsen, M.N., Christensen, J.H., Kruhoffer, M., Palmfeldt, J., Corydon, T.J., Gregersen, N., Ang, D., et al. (2008). The Hsp60-(p.V98I) mutation associated with hereditary spastic paraplegia SPG13 compromises chaperonin function both in vitro and in vivo. J Biol Chem 283, 15694–15700. 10.1074/jbc.M800548200.

58. Carlsen, J., Cömert, C., Bross, P., and Palmfeldt, J. (2020). Optimized High-Contrast Brightfield Microscopy Application for Noninvasive Proliferation Assays of Human Cell Cultures. Assay Drug Dev. Technol. 18, 215–225. 10.1089/adt.2020.981.

59. Cömert, C., Fernandez-Guerra, P., and Bross, P. (2019). A Cell Model for HSP60 Deficiencies: Modeling Different Levels of Chaperonopathies Leading to Oxidative Stress and Mitochondrial Dysfunction. Methods in molecular biology (Clifton, N.J.) 1873, 225–239. 10.1007/978-1-4939-8820-4_14.

60. Harboe, M., Torvund-Jensen, J., Kjaer-Sorensen, K., and Laursen, L.S. (2018). Ephrin-A1-EphA4 signaling negatively regulates myelination in the central nervous system. Glia 66, 934–950. 10.1002/glia.23293.

61. Labun, K., Montague, T.G., Gagnon, J.A., Thyme, S.B., and Valen, E. (2016). CHOPCHOP v2: a web tool for the next generation of CRISPR genome engineering. Nucleic Acids Res 44, W272–276. 10.1093/nar/gkw398.

62. Brinkman, E.K., Chen, T., Amendola, M., and van Steensel, B. (2014). Easy quantitative assessment of genome editing by sequence trace decomposition. Nucleic Acids Res 42, e168. 10.1093/nar/gku936.

63. Paternoster, V., Svanborg, M., Edhager, A.V., Rajkumar, A.P., Eickhardt, E.A., Pallesen, J., Grove, J., Qvist, P., Fryland, T., Wegener, G., et al. (2019). Brain proteome changes in female Brd1(+/-) mice unmask dendritic spine pathology and show enrichment for schizophrenia risk. Neurobiol Dis 124, 479–488. 10.1016/j.nbd.2018.12.011.

64. Patro, R., Duggal, G., Love, M.I., Irizarry, R.A., and Kingsford, C. (2017). Salmon provides fast and bias-aware quantification of transcript expression. Nat Methods 14, 417–419. 10.1038/nmeth.4197.

65. Love, M.I., Soneson, C., Hickey, P.F., Johnson, L.K., Pierce, N.T., Shepherd, L., Morgan, M., and Patro, R. (2020). Tximeta: Reference sequence checksums for provenance identification in RNA-seq. PLoS Comp. Biol. 16, e1007664. 10.1371/journal.pcbi.1007664.

66. Love, M.I., Huber, W., and Anders, S. (2014). Moderated estimation of fold change and dispersion for RNA-seq data with DESeq2. Genome Biol 15, 550. 10.1186/s13059-014-0550-8.

67. Birkler, R.I., Stottrup, N.B., Hermannson, S., Nielsen, T.T., Gregersen, N., Botker, H.E., Andreasen, M.F., and Johannsen, M. (2010). A UPLC-MS/MS application for profiling of intermediary energy metabolites in microdialysis samples--a method for high-throughput. J Pharm Biomed Anal 53, 983–990. 10.1016/j.jpba.2010.06.005.

68. Smith, C.A., Want, E.J., O’Maille, G., Abagyan, R., and Siuzdak, G. (2006). XCMS: processing mass spectrometry data for metabolite profiling using nonlinear peak alignment, matching, and identification. Anal Chem 78, 779–787. 10.1021/ac051437y.

69. Durinck, S., Spellman, P.T., Birney, E., and Huber, W. (2009). Mapping identifiers for the integration of genomic datasets with the R/Bioconductor package biomaRt. Nat Protoc 4, 1184–1191. 10.1038/nprot.2009.97.

